# TSvelo: Comprehensive RNA velocity by modeling the cascade of gene regulation, transcription and splicing

**DOI:** 10.1101/2024.12.24.630058

**Authors:** Jiachen Li, Zhe Wang, Hong-Bin Shen, Ye Yuan

**Affiliations:** State Key Laboratory of Biopharmaceutical Preparation and Delivery, Institute of Process Engineering, Chinese Academy of Sciences, Beijing 100190, China; Institute of Image Processing and Pattern Recognition, Shanghai Jiao Tong University, and Key Laboratory of System Control and Information Processing, Ministry of Education of China, Shanghai, 200240, China

## Abstract

RNA velocity approaches fit gene dynamics and infer cell fate by modeling the splicing process using single-cell RNA sequencing (scRNA-seq) data. However, due to short time scale of splicing, high noise and large complexity of data, existing RNA velocity methods often fail to precisely capture the complex velocity dynamics for individual gene and single cell, which makes its downstream analysis less reliable and less robust. We propose **TSvelo**, a comprehensive RNA **velo**city mathematics framework that can model the cascade of gene regulation, **T**ranscription and **S**plicing using highly interpretable neural Ordinary Differential Equations (ODEs). TSvelo can precisely capture the transcription-unspliced-spliced 3D dynamics of all genes simultaneously, infer unified latent time shared by genes within single cell, and be applied to multi-lineage datasets. Experiments on six scRNA-seq datasets, including two multi-lineage datasets, demonstrate TSvelo’s superiority.

## Introduction

Single-cell RNA sequencing (scRNA-seq) enables the detailed exploration of gene expression at the individual cell level. To move beyond static snapshots and capture the dynamic behavior of cells over time, several trajectory inference (TI) methods have been developed, such as PAGA^1^, Monocle^2^, Slingshot^3^ and Palantir^4^. These methods typically estimate pseudotime using diffusion processes and require prior annotation of initial cells. In contrast, RNA velocity^5^ offers a more interpretable approach by modeling the time derivative of gene expression, linking unspliced (immature) and spliced (mature) mRNA levels through Ordinary Differential Equations (ODEs).

Several RNA velocity methods have been proposed to capture splicing dynamics. The first approach, Velocyto^5^, uses least squares solutions to estimate parameters under the assumption of steady-state kinetics. ScVelo^6^ improves upon this by employing an Expectation-Maximization (EM) approach for better fitting splicing kinetics. UniTVelo^7^ propose a top-down idea that directly models the spliced RNA levels with a time-dependent function. More recently, generative models such as VeloVI^8^, veloVAE^9^, Pyrovelocity^10^ and BayVel^11^ have been introduced, utilizing Bayesian frameworks to estimate RNA velocity. To handle multi-lineage datasets, methods like CellDancer^12^, DeepVelo^13^ and LatentVelo^14^ extend RNA velocity by modeling local dynamics with neural networks, rather than assuming a globally constant transcriptional rate. Apart from using unspliced/spliced data, Dynamo^15^ enhances RNA velocity further by incorporating labeled RNA-seq data. The advent of single-cell multi-omics^16,17^ technologies has allowed RNA velocity analysis to extend to protein abundance (e.g., protaccel^18^) and single-cell ATAC-seq datasets (e.g., MultiVelo^19^). Additionally, methods like DeepCycle^20^ and VeloCycle^21^ have been developed to focus on cell cycle processes, with specialized modules for capturing periodic signals. STT^22^ and SIRV^23^ proposed the idea on extending RNA velocity analysis to spatial transcriptomics.

Although RNA velocity theory has significantly advanced the inference of single-cell trajectories, pseudotime, and gene regulation^24^, several challenges persist for current RNA velocity models. Firstly, RNA velocity models primarily infer cell fate based on phase portrait fitting of unspliced and spliced dynamics for each gene. However, they often fail to capture the correct phase portrait for most genes, due to the sparsity and noise in unspliced and spliced mRNA abundance for individual genes, the short time scale of the splicing process, and the mixing of cells from different types on the phase portrait^25-27^. Secondly, The majority of existing RNA velocity models treat each gene independently and fail to incorporate the underlying regulatory interactions^28^. Although some approaches (e.g. TFvelo^27^, PHOENIX^29^, scKINETICS^30^ and scPN^31^) have constructed ODE models to capture gene dynamics by integrating regulation information, these methods overlook the splicing signal so that they cannot jointly model the transcription and splicing dynamics into one unified form. Thirdly, classical RNA velocity approaches, such as scVelo, use interpretable parameters in constructing dynamic model for single gene. By contrast, to model flexible transcriptional rates or integrate multiple genes, several recent methods employ latent space embeddings or neural network-based encoders^12^. It makes the model parameters less interpretable in detailed gene level, which is crucial for understanding the underlying biological mechanisms. Fourthly, multi-lineage tasks remain a significant challenge for current RNA velocity models due to the complexity in large scale scRNA-Seq datasets.

To address the challenges outlined above, we propose TSvelo, a method that intergrates gene regulation, transcription and splicing of all genes into a single ODE model, whose parameters are highly interpretable. Using a high-dimentional Neural ODE solver^32,33^, TSvelo directly learns the global latent time without the need to separately learn gene-specific latent times. By leveraging both unspliced and spliced scRNA-seq data, along with gene regulatory knowledge from TF-target databases, TSvelo iteratively optimizes the parameters in the ODE model and the unified latent time using Expectation-Maximization (EM) algorithm. Experiments on six scRNA-seq datasets, including two multi-lineage datasets, demonstrate that TSvelo outperforms existing methods in modeling gene dynamics, inferring cell fates, and is also effective in analyzing multi-lineage datasets.

## Results

### Estimate RNA velocity with TSVelo

We present TSvelo, a method for jointly estimating RNA velocity across high-dimentional genes by integrating both transcriptional regulation and splicing information. Existing RNA velocity models predominantly focus on modeling the splicing process and typically operate in a gene-wise manner. In contrast, TSvelo explicitly models the full cascade of regulation, transcription, and splicing, while capturing their coordinated dynamics across all genes simultaneously. A systematic comparison between TSvelo and representative RNA velocity methods is provided in **Table S1** at Supplementary Information. In TSvelo, the unspliced and spliced RNA abundances are initially preprocessed for velocity gene selection and pseudotime initialization **(Fig. 1a)**. See **Methods** for details about preprocessing. Next to model the velocity gene *g*’s expression dynamics, we suppose *u*_*g*_ and *s*_*g*_ are the abundance of unspliced and spliced RNA, and *α*_*g*_ (*t*), *β*_*g*_, *γ*_*g*_ are the transcription 105 rate, splicing rate and degradation rate, respectively. The dynamics is modeled as:

**Figure 1.**
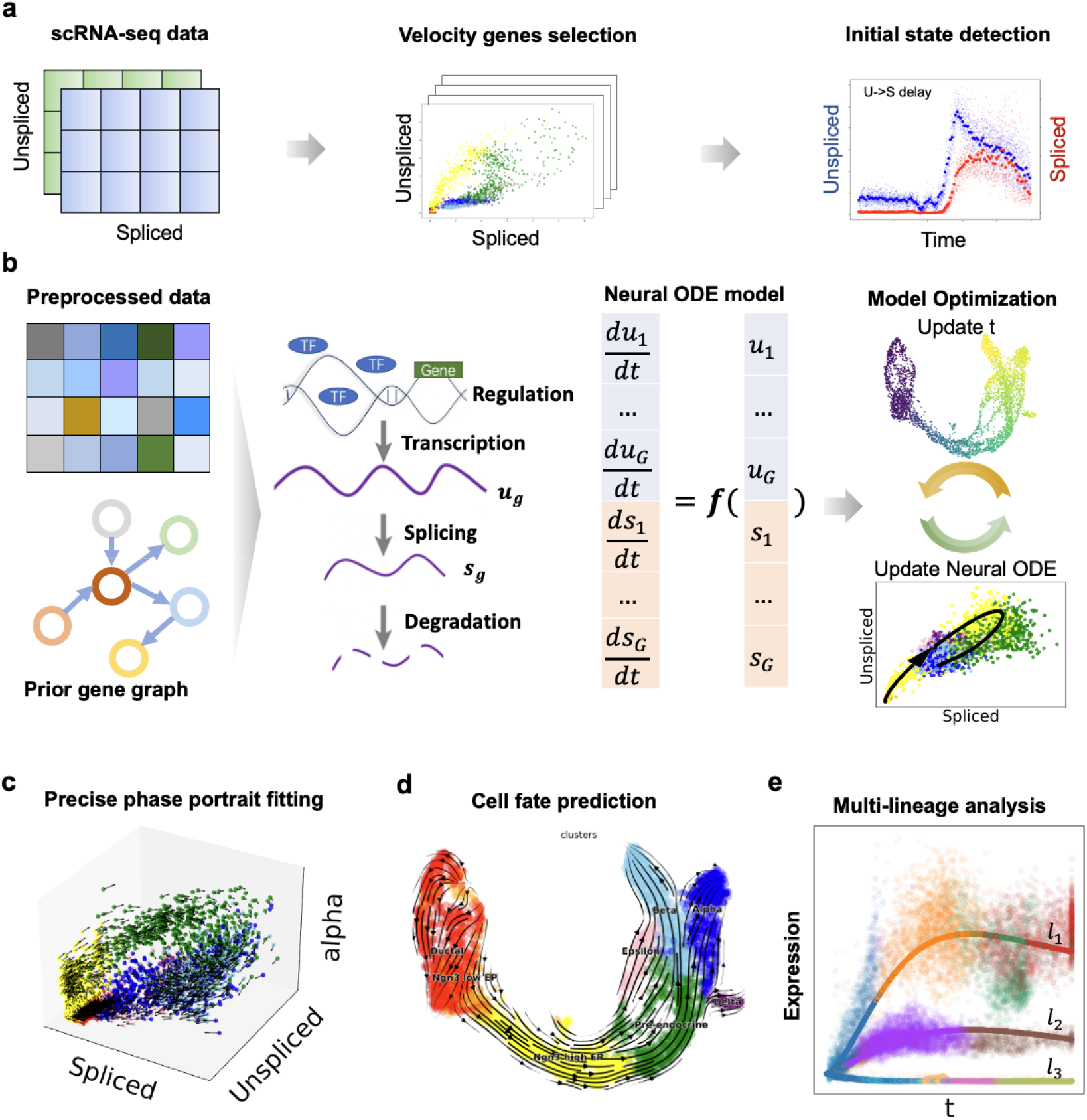
The framework of TSvelo. **(a)** The preprocessing strategy in TSvelo, including velocity genes selection and initial state detection. **(b)** The Neural ODE model and its optimization in TSvelo, where the parameters and latent time is optimized iteratively. (**c-e)** The downstream application tasks of TSvelo, including precise transcription-unspliced-spliced 3D phase portrait fitting (c), cell fate prediction using predicted RNA velocity (d) and gene expression pattern analysis for multi-lineage dataset (e).

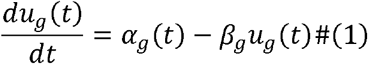

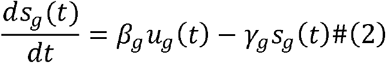

Furthermore, we assume that the gene and cell-specific transcriptional rate *α*_*g*_ (*t*) is influence by the expression of Transcriptional Factors (TFs). Considering the wide usage and interpretability of linear models for gene-relations in previous studies^27,34-37^, we model *α*_*g*_ (*t*) as

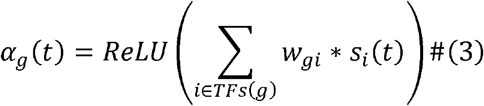

*ReLU* denotes the Rectified Linear Unit activation function, defined as *ReLu* = *x max*(*0,x)*.the term *w*_*gi*_ represents the regulatory weight of TF *i* on the target gene *g. TFs*(*g*) refers to the TFs that could regulated the target gene *g*,which are selected according to the ChEA and ENCODE TF-target database. In order to include more TFs in the model, we also reserve those TFs that are not selected as velocity genes, and directly model the dynamics between transcription and spliced RNA without using the unspliced abundance. Finally, we combine the dynamic models on all selected genes into the Ordinary Differential Equation (ODE) matrix form **(Fig. 1b**), which enables directly inferring a unified latent time for each cell using its all genes. See **Methods** for details.

TSvelo employs an EM framework to iteratively optimize both latent time and the parameters in ODE. The global pseudotime is assigned to each cell through grid search. Due to the difficulty in calculating analytical solutions of or. TSvelo adopted a Neural ODE for estimating those parameters of transcription rate, splicing rate and degradation rate. Next, TSvelo can model the gene dynamics of both transcription, unsplicing and splicing processes **(Fig. 1c)**, predict cell states using both pseudotime and velocity stream analysis **(Fig. 1d)**, and is also applicable to multi-lineage tasks for analyzing expression patterns across different lineages in large complex scRNA-Seq datasets **(Fig. 1e)**. We further validated TSvelo on simulated datasets and observed consistent improvements in velocity estimation and trajectory reconstruction across a range of settings **(Fig. S1** and **Fig. S2** in the Supplementary Information)

### TSvelo can model 3D gene dynamics and predict cell fate on pancreas dataset

To validate the TSvelo model, we applied it to the pancreas scRNA-seq dataset^5^, which is widely used in RNA velocity studies to model cell differentiation from ductal to endocrine cells. Using the predicted RNA velocity, TSvelo can generate velocity stream plot using the functions provided in scVelo as well. Both the pseudotime *t* **(Fig. 2a)** and the velocity stream plot **(Fig. 2b)** effectively capture the cell differentiation process. We quantitively compare TSvelo and baseline approaches, including scVelo^6^, dynamo^15^, UniTVelo^7^, cellDancer^12^ and TFvelo^27^.

**Figure 2.**
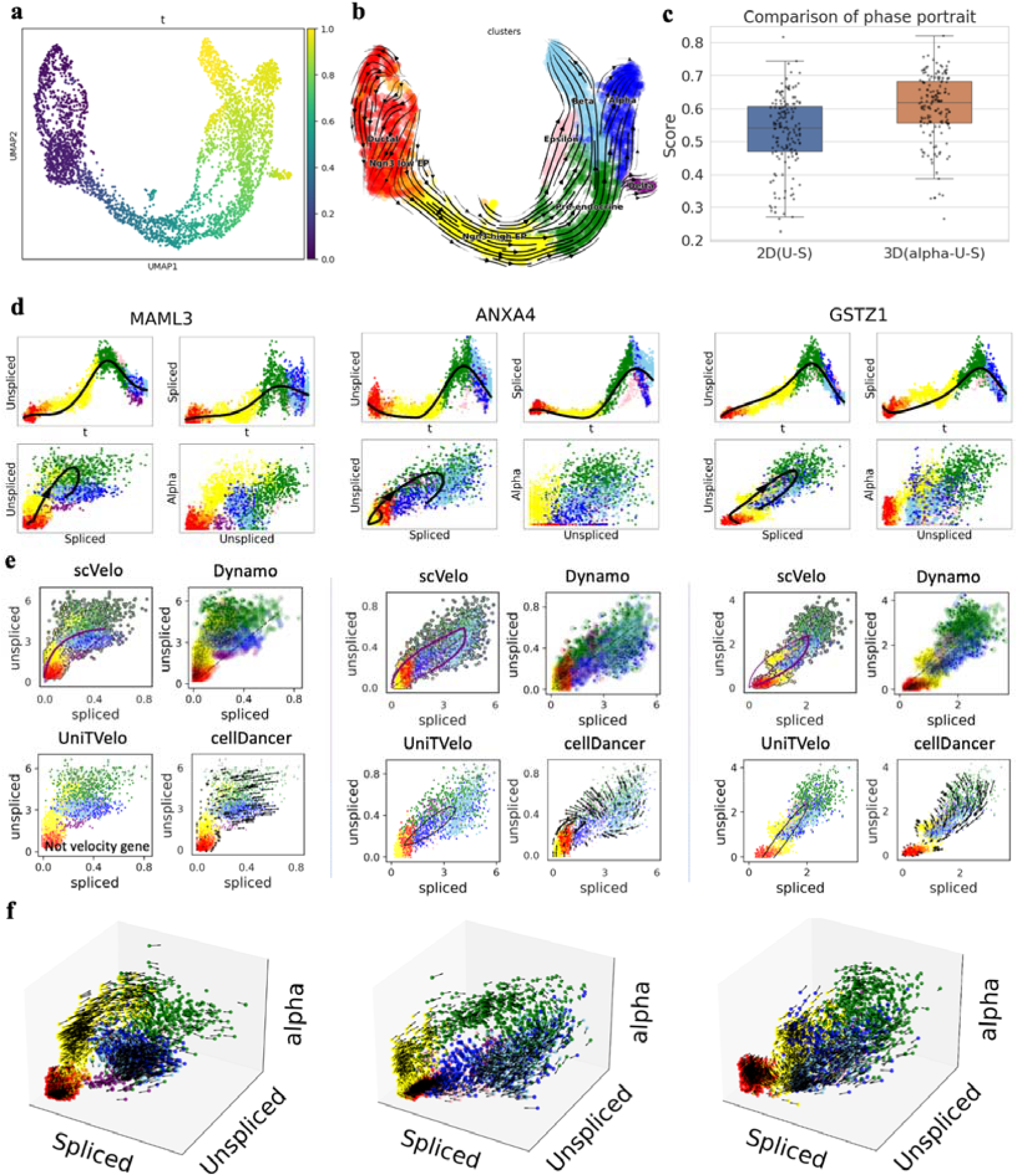
Results on pancreas dataset. **(a)** The pseudotime learned with TSvelo. **(b)** The Stream-plot for visualization the RNA velocity inferred by TSvelo. **(c)** A quantitative comparison of cell-state separability between the 3D phase portrait used in TSvelo and the traditional 2D phase portrait. The down, central and up hinges correspond to the first quartile, median value and third quartile, respectively. The whiskers extend to 1.5× the interquartile range of the distribution from the hinge. 3,696 samples are included in the boxplot for each method. **(d)** The dynamics fitting on MAML3, ANXA4 and GSTZ1. For each gene, four plots are displayed in a 2×2 layout: the u-t, s-t, u-s, and alpha-u plots. (**e)** The unspliced-spliced phase portrait fitting on MAML3, ANXA4 and GSTZ1 obtained by four baseline RNA velocity approaches, including scVelo, Dynamo, UniTVelo and cellDancer. **(f)** The dynamics fitting in transcription-unspliced-spliced 3D phase portrait for MAML3, ANXA4 and GSTZ1.

In the conventional 2D unspliced–spliced phase portrait, cells from different clusters often overlap, limiting separability. By introducing the latent variable *α*, the representation is extended to a 3D space, which helps disentangle these mixed states and reveals a clearer phase structure. As shown in **Fig. 2c**, the 3D representation achieves consistently higher kNN classification accuracy for cell-state separation than the 2D u–s embedding (one-sided Mann–Whitney U test, p = 4.37 x 10 □ ^1^□ · Detials are provided in **Methods)**. These results indicate that the 3D phase portrait provides improved separation of cell states and a stronger foundation for modeling the underlying dynamics along cell trajectories.

Additional quantitative evaluation is provided in **Fig. S4** at Supplementary Information. TSvelo achieves the highest median velocity consistency, which demonstrates that the high-dimensional velocity vectors learned by TSvelo are mostly coherent within neighbor cells. TSvelo also achieves the highest median in-cluster coherence and cross-boundary correctness, which validates that TSvelo best fit the differentiation process within these cell types according to the ground truth annotation.

Next, we show the dynamics for individual genes and demonstrate how the TSvelo model aids gene dynamics analysis by incorporating both transcriptional and splicing information. MAML3, ANXA4 and GSTZ1 are selected as examples because their unspliced–spliced 2D phase portrait exhibits mixed or overlapping patterns that are difficult to model using conventional RNA velocity approaches. **Fig. 2d** present the results of the TSvelo model on these genes, and **Fig. 2e** present the results of baseline approaches. Many previous approaches assume that the unspliced-spliced phase portrait exhibits an almond-shaped distribution^38^which may not actually hold in the data. For MAML3, while the unspliced-spliced distribution predominantly follows the almond-shaped pattern, some cells, specifically Alpha cells (blue) and Beta cells (light blue), overlap with Ngn3 high EP cells (yellow), indicating that these cell types cannot be distinctly separated using only the unspliced-spliced 2D phase portrait. Nevertheless, TSvelo can infer transcription representation from the expression of multiple transcription factors, which enables the model to distinguish these cell types **(Fig. 2d** and **Fig. 2f)**. ANXA4 shows higher expression in Ductal cells (in red) compared to Ngn3 low EP cells (in orange), which mean its expression pattern exhibits an initial decrease followed by an increase. Such dynamics are not easily captured in the conventional unspliced–spliced phase portrait used by previous approaches, as many baseline methods implicitly assume a decreasing–then–increasing expression pattern. By comparison, TSvelo can still fit such expression pattern by using additional information from the 3D phase portrait. The visualization of dynamics fitting on more genes are provided at **Fig. S5** and **Fig. S6 in Supplementary Information**.

### TSvelo can better predict RNA velocity on gastrulation erythroid dataset

We applied TSvelo to the gastrulation erythroid dataset, which is derived from the transcriptional profile of mouse embryos ^39^ and has been used in previous RNA velocity studies. This dataset primarily describes the differentiation process from blood progenitors to erythroid cells. First, using Gene Ontology (GO) term enrichment analysis **(Fig. 3a**), we find that the velocity genes selected through TSvelo’s preprocessing strategies are mostly enriched in the to the erythropoiesis-related process, providing a preliminary validation of the biological plausibility of the inferred velocities.

**Figure 3.**
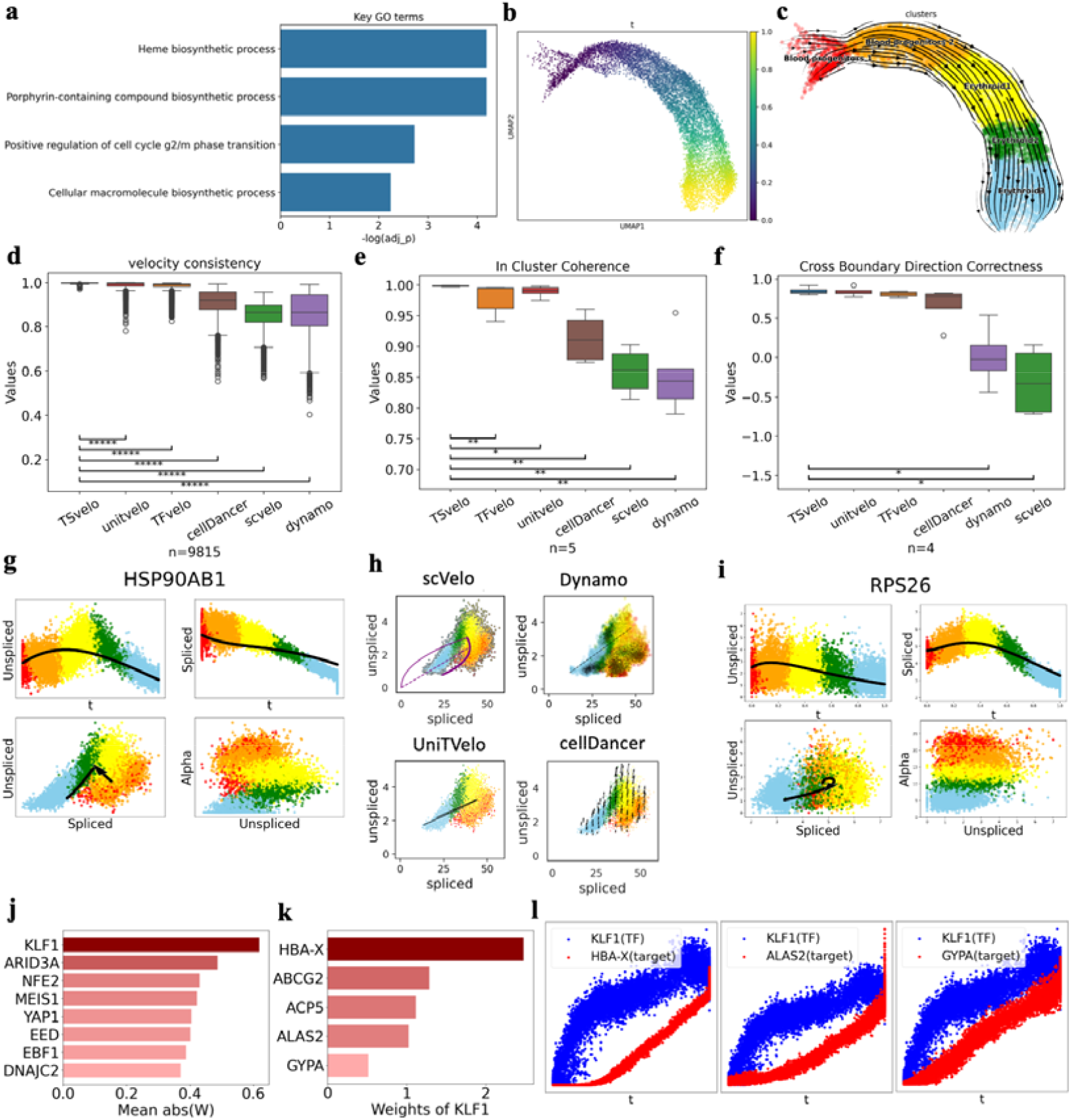
Results on gastrulation erythroid dataset. **(a)** The GO terms which are mostly enriched in the selected velocity genes of TSvelo. **(b)** The pseudotime learned with TSvelo. **(c)** The Stream-plot for visualization the RNA velocity inferred by TSvelo. **(d-f)** The quantitative comparison between TSvelo and multiple baseline approaches in terms of velocity consistency (d), in-cluster coherence (e), and cross boundary direction correctness (f). The down, central and up hinges correspond to the first quartile, median value and third quartile, respectively. The whiskers extend to 1.5× the interquartile range of the distribution from the hinge. 9,815 samples are included in the boxplot for each method. **(g)** The dynamics fitting on HSP90AB1 obtained by TSvelo. Four plots are displayed in a 2×2 layout: the u-t, s-t, u-s, and alpha-u plots, where u, a and alpha mean the abundance of unspliced mRNA, the abundance of spliced mRNA, and the learned transcriptional representation, respectively. **(i)** The phase portrait fitting on HSP90AB1 obtained by baseline approaches. **(i)** The dynamics fitting on RPS26 obtained by TSvelo. **(j)** The TFs with the highest ranked weights as identified by TSvelo. **(k)** Weights for KLF1’s targets, with the highest absolute weight. **(l)** Temporal dynamics of KLF1 and its target genes with the highest weights along pseudotime, which includes HBA-X, ALAS2 and GYPA.

After applying TSvelo, both the inferred pseudotime **(Fig. 3b)** and the velocity stream plot **(Fig. 3c)** are compatible with current biological knowledge about the cell differentiation process. We also compare TSvelo with previous approaches. The results, shown in **Fig. 3d-f**, demonstrate that TSvelo achieves the highest velocity consistency, the highest in-cluster coherence, and the highest cross-boundary correctness, showing that the high-dimensional dynamics predicted by TSvelo best capture the underlying biological processes.

We next analyze TSvelo modeling at the gene level in **Fig. 3g-i**. The 3D phase portrait again provides a better basis for dynamic fitting, as it more effectively separates cells from different clusters than the traditional 2D phase portrait **(Fig. S7)**. For genes exhibiting higher noise or more complex patterns, TSvelo demonstrates its robustness in dynamic fitting by integrating both transcriptional and splicing signals. For HSP90AB1, which exhibits a counter-clockwise pattern in the unspliced–spliced phase portrait **(Fig. 3g)**, in contrast to the clockwise dynamics typically assumed by most baseline approaches, it is difficult for previous methods to capture this behavior **(Fig. 3h)**, whereas TSvelo can still faithfully model such patterns. For genes such as RPS26, which have critical roles in the development in blood progenitors to erythroid^40^, the unspliced-spliced data is so noisy that cells of different types overlap in phase portrait. TSvelo can still captures the gene dynamics and reveals differences in transcription rates (Alpha) across cell types **(Fig. 3i)**. In contrast, methods that rely solely on unspliced-spliced data from individual genes, such as scVelo, fail to accurately fit these dynamics and can not identify them as velocity genes, which further validates TSvelo’s performance in complex scenarios. The visualization of dynamics fitting on more genes are provided at **Fig. S8** in **Supplementary Information**.

From the TF-target weight matrices learned by TSvelo, we can extract key regulatory relationships. First, we calculate the mean absolute weight of each TF across its interactions with target genes. The TFs with the highest mean absolute weights are ranked and presented in **Fig. 3j**. Notably, KLF1, a critical TF in blood progenitors^41^, is assigned the highest weight. We then examine the target genes of KLF1, highlighting those with the highest weights **(Fig. 3k)**, among which HBA-X^42^, ALAS2^43^, and GYPA^44^ have been previously identified as key genes in erythroid differentiation and are known to be regulated by KLF1. A time-delay pattern is also observed between KLF1 and its target genes **(Fig. 3l)**, indicating that KLF1 expression increases first, followed by a corresponding upregulation of its target genes.

### TSvelo can gene dynamics well and predict cell fate on mouse brain data

We apply TSvelo to a 10x multi-omics dataset from the embryonic mouse brain, which includes both Assay for Transposase-Accessible Chromatin with sequencing (ATAC-Seq) ^45^ and scRNA-seq data. This dataset was previously used in the Multivelo study, which introduced an RNA velocity model designed for multi-omics datasets to capture the dynamics between chromatin accessibility and unspliced mRNA^20^. As introduced by previous study^19^, Radial Glia (RG) cells located in the outer subventricular zone give rise to neurons, astrocytes, and oligodendrocytes. During cortical development, neurons follow an inside-out layering pattern in which earlier-born neurons populate the deep cortical layers, whereas later-born neurons migrate past them to occupy more superficial layers^46^. Additionally, RG cells are capable of producing intermediate progenitor cells (IPCs), which serve as neural stem cells and generate a variety of mature excitatory neurons within the cortical layers.

Multivelo is chosen as the baseline on this study since it connects the regulation signal and RNA velocity in ATAC-unspliced-spliced space. TSvelo uses the same data processed be Multivelo for a fair comparison. The velocity stream obtained by TSvelo and Multivelo are shown in **Fig. 4a** and **Fig. 4b**, respectively. TSvelo produces velocity patterns that appear more consistent with expected differentiation trends, especially for the RG (in red) to IPCs (in brown) process. The pseudotime inferred by TSvelo is also coherent with the whole cell development process **(Fig. 4c)**.

**Figure 4.**
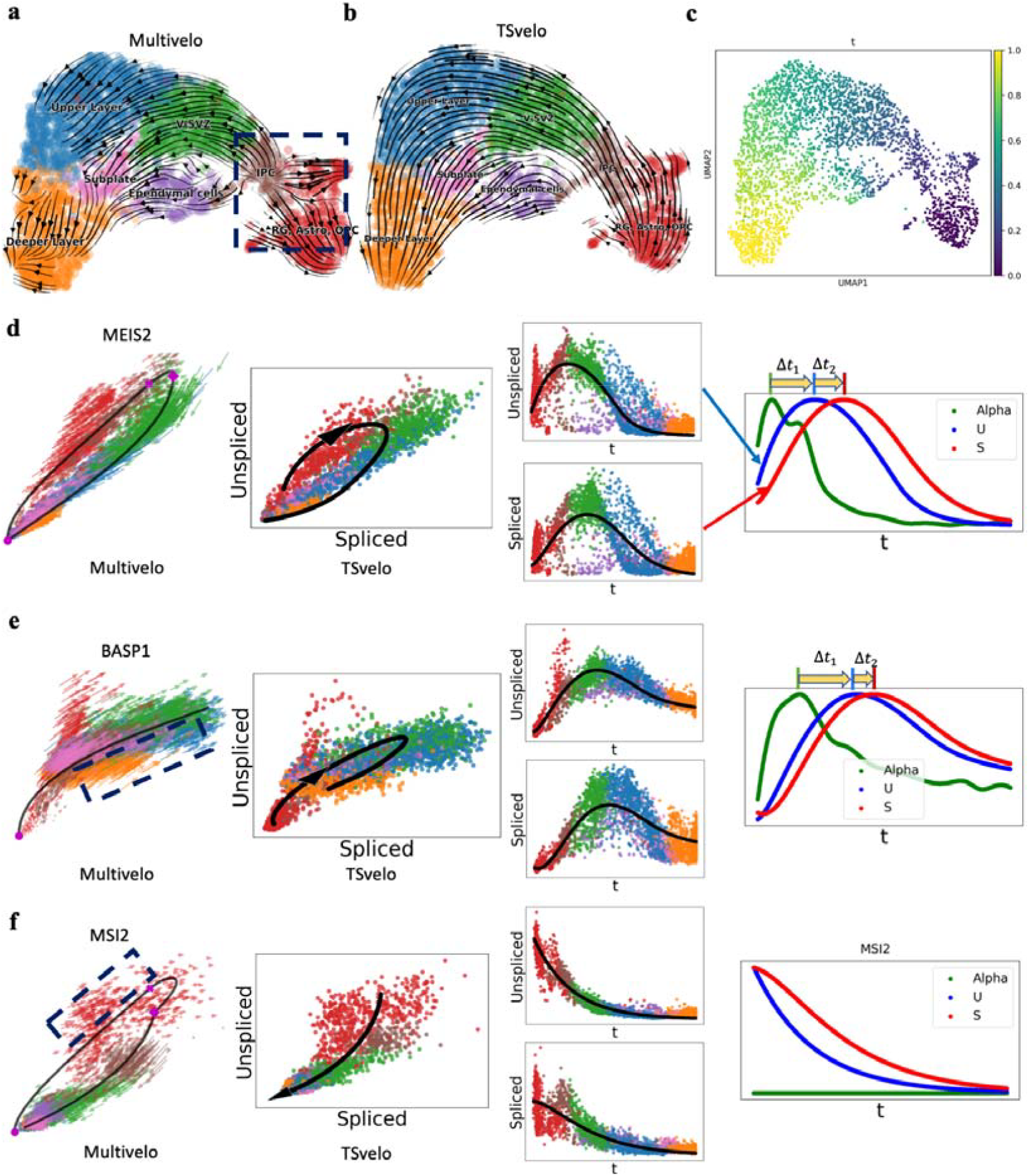
Results on mouse brain dataset. **(a)** The velocity stream inferred by Multivelo. **(b)** The Stream-plot for visualization the RNA velocity inferred by TSvelo. (c)The pseudotime learned with TSvelo. **(d-f)** The dynamics fitting on gene MEIS2 (d), BASP1 (e) and MSI2 (f) by both Multivelo and TSvelo. In each panel, the leftmost plot shows the phase portrait fitting of Multivelo, and the next two columns show TSvelo’s results. The rightmost plot shows the learned transcriptional rate (in green), unspliced abundance (in blue), and spliced abundance (in red) along the pseudotime. Since the transcriptional rate is calculated for each individual cell, we apply a Generalized Additive Model (GAM) to transcriptional representation across cells along the pseudotime and present GAM-fitting results to better visualize its trends in these plots.

We next further analyze the details in the phase portrait fitting. On MEIS2 **(Fig. 4d)**, both Multivelo and TSvelo successfully models its dynamics, which follows a clear pattern on the unspliced-spliced phase portrait. However, due to the high noise and sparsity inherent in ATAC-seq data, Multivelo encounters difficulty on genes that show significant overlap between different cell types in the unspliced–spliced phase portrait. For example, in the case of BASP1 **(Fig. 4e)**, Multivelo incorrectly models their expression patterns as monotonically increasing, failing to capture the true dynamics, which involve initial upregulation followed by downregulation. In contrast, TSvelo’s prediction consistents with known biological processes and identifies the delay from transcription to the unspliced mRNA and also the delay from unspliced to spliced mRNA (The rightmost plots in **Fig. 4e)**. MSI2, which has been reported to be highly expressed in neural stem/progenitor cells^47^, is more accurately modeled by TSvelo, exhibiting a decreasing expression pattern during the differentiation process. By comparison, Multivelo fails to capture the correct trend at the initial stage **(Fig. 4f)**. We also show the learned transcriptional rate *α* as well as the predicted dynamics of unspliced and spliced RNA along pseudotime *t*, which clearly illustrate the time delays between transcription to unspliced and unspliced to spliced RNAs. The visualization of dynamics fitting on more genes are provided at **Fig. S9** in **Supplementary Information**. Given that transcriptional regulation involves the binding of TF and chromatin accessibility as measured by ATAC, TSvelo shows that it is possible to model transcriptional signals using only scRNA-Seq data.

### TSvelo can predict cell fate and model lineage-specific gene dynamics for multi-lineage task

Given that multi-lineage differentiation is a common phenomenon in larger and larger scRNA-seq datasets, developing the RNA velocity model that can handle such complexity is essential. Because TSvelo demonstrates robust performance across various tasks, its applicability can be extended to more complex situations, such as scRNA-seq datasets where cells differentiate into multiple fates. We apply TSvelo to a multi-lineage dentate gyrus scRNA-seq dataset, which captures the differentiation process from neural blast cells to various cell types^48^. During preprocessing, the velocity genes selected by TSvelo are enriched in GO terms related to neural development, such as axonogenesis (GO:0007409), axon guidance (GO:0007411) and axon development (GO:0061564) **(Fig. 5a)**, which aligns with the biological processes described by the dataset.

**Figure 5.**
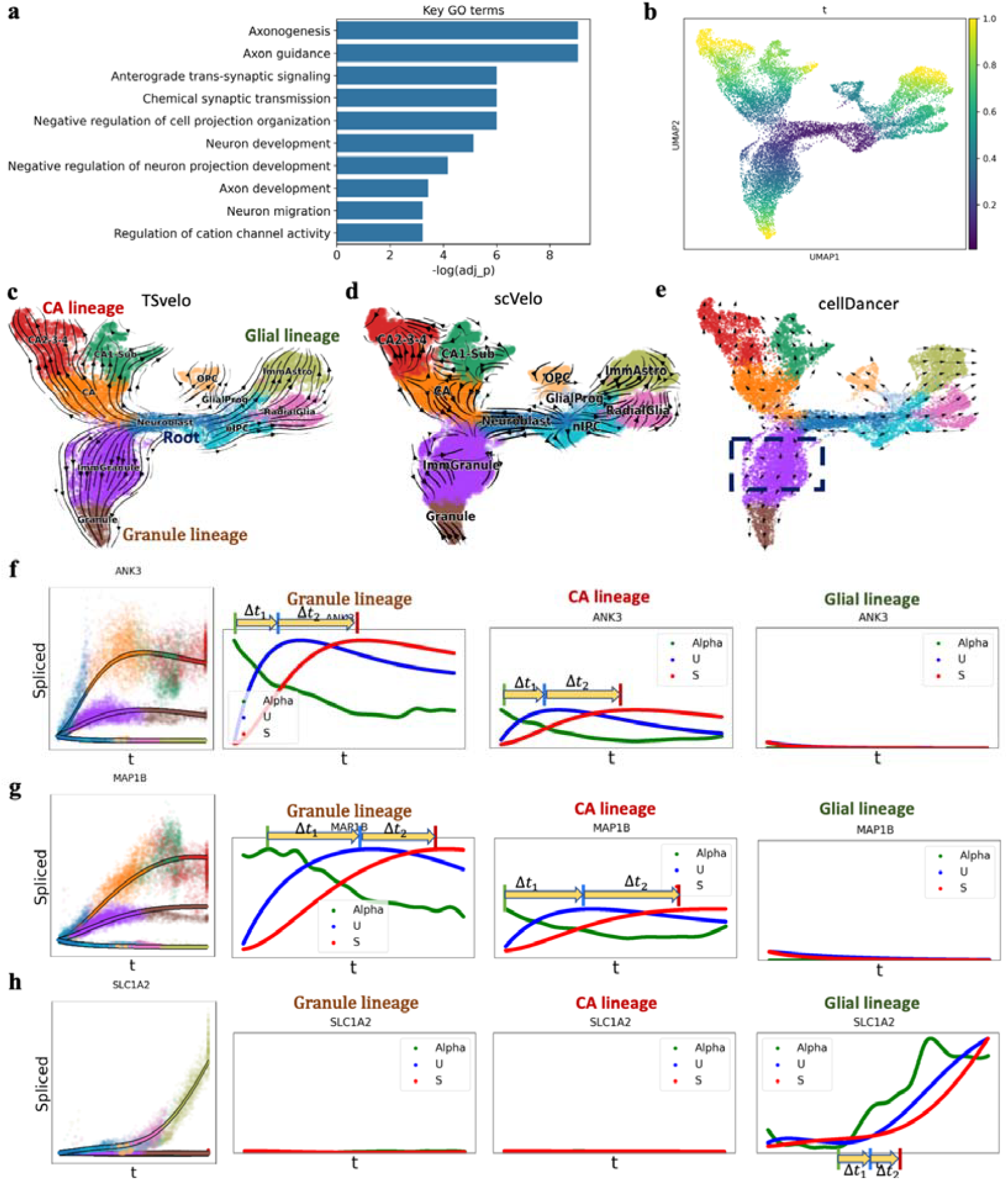
Results on the multi-lineage dentate gyrus dataset. **(a)** The GO terms enriched in the selected velocity genes of TSvelo. **(b)** The pseudotime learned with TSvelo. **(c)** The Stream-plot for visualization the RNA velocity inferred by TSvelo. Three lineages are detected, which are Granule lineage, CA lineage and glial lineage. **(d)**The velocity stream inferred by scVelo. **(e)** The velocity stream inferred by cellDancer. **(f-h)** The dynamics modeling of TSvelo on three axonogenesis-related genes, ANK3 (f), MAP1B (g) and SLC1A2 (h). In the leftmost plot of each panel, the lines represent the predicted spliced abundance across all lineages, with the color indicating the cell types most associated with each pseudotime point along the corresponding lineage. Additionally, expression data for each lineage are shown as translucent points. The remaining plots in each panel display the dynamics of the learned transcriptional rate (in green), unspliced abundance (in blue), and spliced abundance (in red) along pseudotime for each lineage. The transcriptional representation in these plots is also processed using GAM fitting.

TSvelo could correctly identify three lineages from this data. The lineage segmentation process is fully automated and does not require prior knowledge about the presence of multiple branches (details are provided at **Methods** and **Fig. S10** in **Supplementary Information)**. After applying the TSvelo model to each lineage and combine the results, the inferred pseudotime and velocity stream plots **(Fig. 5b** and **Fig. 5c)** align with the ground truth differentiation trajectory well, which begins with neural blast cells. We also compare TSvelo to baseline methods, including scVelo and cellDancer. Notably, cellDancer is designed to handle such multi-lineage data by modeling cell- and gene-specific transcriptional rates, splicing rates, and degradation rates using neural networks. As shown in **Fig. 5d** and **Fig. 5e**, scVelo struggles with this multi-lineage dataset, and cellDancer fails to correctly capture the trajectory of immature granule cells (colored in purple).

Next, we analyze gene expression patterns across different lineages. TSvelo provides inferred dynamics along each lineage, revealing that the expression of ANK3 in the granule and Cornu Ammonis (CA) lineages follows an increasing and then decreasing pattern. In contrast, the expression of ANK3 in the glial lineage decreases to zero **(Fig. Sf)**. The similar pattern is also observed for other genes, such as MAP1B **(Fig. 5g)**. Both ANK3^49^ and MAP1B^50^ are associated with GO terms axonogenesis and axon guidance. This observation is consistent with the fact that the granule and Cornu Ammonis^51^ predominantly consist of neurons, where axons are a critical component. In contrast, glial cells, such as astrocytes, typically lack axons, providing a potential explanation for these expression patterns. SLC1A2 exhibits a distinct expression pattern, showing a significant increase in the glial lineage and a decrease in the granule and CA lineages. This pattern aligns with the observation that astrocytes are the primary cell type expressing SLC1A2^52^. Details about dynamics fitting for genes on each lineage are provided at **Fig. S11 in Supplementary Information**.

### TSvelo can predict cell fate and model lineage-specific gene dynamics on Larry dataset

The Lineage and RNA Recovery (LARRY) method utilizes barcoded hematopoietic cells to trace both cell lineage and gene expression over time. LARRY has been successfully employed to track the in vitro differentiation of human blood cells, accurately capturing lineage trajectories and cell fates^53^.

On this LARRY dataset which encompass a total of 49,302 cells on multiple lineages, TSvelo effectively detects the initial Leiden clusters based on the unspliced-to-spliced delay **(Fig. 6a** and **Fig. 6b)** and separate its lineages (Details are provided at the **Lineage segmentation and pseudotime initialization** section in **Methods)**. Furthermore, **Fig. 6c** and **Fig. 6d** show that the pseudotime and velocity stream inferred by TSvelo can capture the differentiation process, progressing from undifferentiated cells to distinct cell fates. The velocity genes selected by TSvelo are significantly enriched in processes related to neutrophil biology, such as neutrophil degranulation (GO:0043312), neutrophil activation in immune response (GO:0002283), and neutrophil-mediated immunity (GO:0002446).

**Fig 6.**
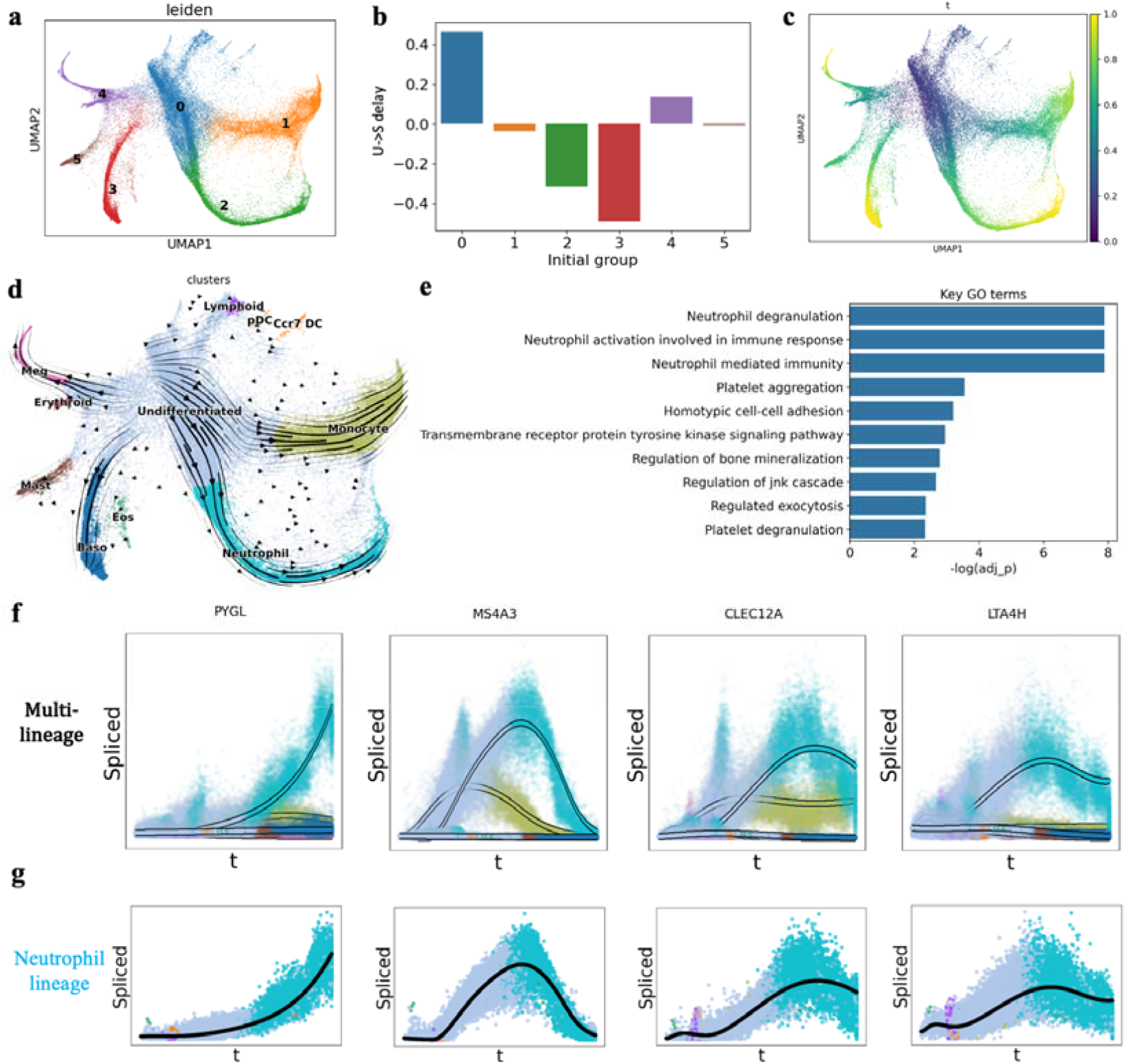
Results on the LARRY dataset. **(a)** The Leiden clustering on LARRY. **(b)** The initial Leiden cluster detection in preprocessing. (**c)** The pseudotime learned with TSvelo. **(d)** The Stream-plot for visualization the RNA velocity inferred by TSvelo. **(e)** The GO terms enriched in the selected velocity genes of TSvelo. **(f)** The dynamics modeling of TSvelo on four genes related to neutrophil development, which are PYGL, MS4A3, CLEC12A and LTA4H. The lines represent the predicted spliced abundance across all lineages, with the color indicating the cell types most strongly associated with each pseudotime point along the corresponding lineage. (g) The dynamics modeling of PYGL, MS4A3, CLEC12A and LTA4H on the neutrophil lineage.

We further analyzed the expression patterns of genes associated with neutrophil degranulation, focusing specifically on the neutrophil lineage. Using four representative genes as examples **(Figs. 6f, g)**, TSvelo can help observe the significant variation in the expression patterns across these genes. PYGL, which has been widely reported as a key gene in neutrophil^54^, is almost exclusively expressed in neutrophil cells and exhibits an increasing expression pattern during neutrophil differentiation. In contrast, MS4A3 was examined across all lineages, and using TSvelo, we observed that its expression initially increases and then decreases in both the neutrophil and monocyte lineages. This pattern is consistent with prior studies identifying MS4A3 as a gene specifically expressed by granulocyte-monocyte progenitors^47^. Similarly, CLEC12A, another gene associated with neutrophil degranulation, shows a comparable expression trajectory during neutrophil differentiation. Previous research has indicated that CLEC12A expression is highest in granulocyte-macrophage progenitors^55^. LTA4H exhibits a pattern similar to that of CLEC12A, suggesting it may play a significant role in the early stages of neutrophil differentiation. These findings further confirm the utility of TSvelo for gene-level analysis in multi-lineage differentiation datasets.

## Discussion

RNA velocity, utilizing unspliced/spliced data, has become a widely adopted concept for predicting cell fate and modeling gene dynamics. While several RNA velocity models have been proposed, most of they are based on phase portrait fitting in the unspliced-spliced space. However, due to the limited information and high noise inherent in the splicing data of individual genes, the short time delays between unspliced and spliced abundance, and the challenges posed by large-scale datasets with complex processes, most genes cannot be accurately modeled by previous RNA velocity methods. This limitation reduces the reliability and robustness of downstream analyses.

Gene expression is a complex biological process within cells, involving multiple regulatory mechanisms from DNA to the matured RNA. In this process, transcription and splicing are two crucial steps to determine the final gene expression. Due to the mathematical complexity, previous methods cannot jointly model gene regulation, transcription and splicing. TSvelo comprehensively models the cascade of the whole process using an interpretable ODE framework, allowing for learning dynamics of all velocity genes simultaneously. TSvelo could accurately model gene dynamics, predict cell fate, detect the key regulatory relations, and handle multi-lineage datasets. Results on six scRNA-seq datasets demonstrate that TSvelo is a valuable approach for RNA velocity modeling (results on pons dataset are provided in **Fig. S3** in **Supplementary Information)**.

One limitation of TSvelo is its reliance on predefined TF–target regulatory priors, which may be incomplete and may not fully capture context-specific regulatory relationships in primary tissues or at single-cell resolution. Second, jointly modeling high-dimensional regulatory interactions and RNA kinetics introduces additional computational overhead compared with some existing approaches, as reflected in runtime and memory analyses **(Table R2, Table R3**, and **Fig. S16)**. Third, accurately capturing cell-cycle transitions remains challenging, since cyclic gene-expression programs may violate the assumption of unidirectional state progression and thereby complicate velocity-based trajectory inference. Finally, like many RNA velocity and trajectory inference methods, TSvelo assumes that the input data reflects underlying dynamic biological processes, highlighting the importance of appropriate preprocessing checks when applying such methods to non-dynamic datasets **(Fig. S17)**.

Looking forward, there are several opportunities to further strengthen the framework. Incorporating context-specific regulatory information, such as single-cell chromatin accessibility data, may improve transcriptional modeling. Moreover, TSvelo currently assumes a constant gene-specific splicing and degradation rates. Since splicing factors also play significant roles in regulating other genes during the splicing process^56^, and there are still complex mechanisms of mRNA degradation^57^, further exploration could enhance velocity’s modelling and bring new biological insights in future.

## Methods

### Preprocessing for scRNA-seq data

The unspliced and spliced RNA abundances are preprocessed with multiple steps, which includes highly variable genes (HVGs) selection, normalization, log transformation, k-nearest neighbor (KNN) smoothing, and clustering. Following the data preprocessing procedures outlined in scVelo^6^, we first select the top 2,000 highly variable genes (HVGs) and normalize their expression profiles by dividing by the total counts in each cell. A nearest-neighbor graph (with 30 neighbors by default) was constructed based on Euclidean distances in principal component analysis space (with 30 principal components by default) on log-transformed gene expression data. Subsequently, we compute the first- and second-order moments (mean and uncentered variance) for each cell across its nearest neighbors. These steps are performed by using scvelo.pp.filter_and_normalize() and scvelo.pp.moments().

### Acquiring Prior Knowledge of Gene Regulatory Relations

To construct a prior gene network for the selected HVGs, we utilize TF-target annotations from the ENCODE^58^ and ChEA^59^ databases. If a regulatory interaction between a transcription factor (TF) and its target gene is identified in either of these two databases, the TF is included in the set of regulators for modeling the dynamic behavior of the target gene. During model optimization, the contribution of each prior regulatory edge is adaptively adjusted, allowing unsupported interactions to be effectively down-weighted. As a result, TSvelo is expected to be relatively robust to false-positive TF–target annotations. By contrast, missing true regulatory interactions are not represented in the candidate regulator set and therefore cannot contribute to the learned regulatory dynamics, making the model potentially more sensitive to false negatives in the prior network. Ablation studies on the selection of TF–target resources **(Fig. S13** and **Fig. S14** at Supplementary Information) verify that integrating multiple TF–target resources can improve performance by increasing regulatory coverage and reducing false negatives in the prior network.

### Velocity genes selection

Previous studies^6^ have demonstrated that only a subset of genes, termed “velocity genes,” can be accurately fitted in the unspliced-spliced phase portrait by RNA velocity models. Inspired by the notion that velocity genes provide high-quality data for dynamic modeling in the phase portrait, we propose to select velocity genes during the preprocessing step. The velocity genes are selected based on the premise that they exhibit clear dynamics in the unspliced-spliced phase portrait, which is necessary for precise modeling splicing dynamics. The selection is based on the similarity between cell-cell neighborhood relationships in UMAP space and those observed in gene-specific unspliced-spliced phase portraits. In detail, we first compute the cell-cell neighborhood graph, *Graph*, using scvelo.pp.neighbors() on all highly variable genes (HVGs). Next, we construct an anndata object *adata*_*g*_ for each gene, which contains only the unspliced and spliced expression data of that gene. We then apply scvelo.pp.neighbors() to each *adata*_*g*_ to calculate the gene-specific neighborhood graph, denoted as *Graph*_*g*_. Subsequently, we compute the similarity between *Graph* and each *adata*_*g*_. The top 100 genes with the highest similarity are selected as velocity genes. These genes are characterized by phase portraits that exhibit a structure similar to that of the UMAP space, thereby enhancing the separation of cells from different types. As a result, the splicing dynamics of these velocity genes are more likely to be captured by RNA velocity models.

### lineages segmentation and pseudotime initialization

Based on the normalized gene expression data, we perform Leiden clustering using scanpy.tl.leiden() with a low resolution (default resolution = 0.1). These Leiden clusters are utilized to identify lineages and determine the differentiation direction for each lineage. Subsequently, we apply PAGA (scanpy.tl.paga()) to assess the connectivity between Leiden groups.

By setting one Leiden group as the initial state, we infer pseudotime using diffusion pseudotime (DPT) with scanpy.tl.dpt(). To detect lineages, we start from the selected initial Leiden cluster and filter the PAGA graph by applying a threshold (default = 0.02). Edges with weights below this threshold are removed, leaving only the shortest path from the initial state to each group. The remaining paths are considered as the detected lineages.

Given a lineage, TSvelo could detect the orientation of differentiation along it, which is based on the fact that unspliced RNA precedes spliced RNA in the expression pattern^60^. For each gene, TSvelo computes the Spearman correlation between unspliced and spliced abundance across different moving steps along the DPT. The time number of moving steps along unspliced to spliced expression, which maximizes the Spearman correlation, is considered the U-to-S delay **(Fig. 1a** shows an example of U-to-S delay with gene EML5). The U-to-S delay for each lineage is calculated as the mean U-to-S delay for all genes on it. And the average U-to-S delay across all lineages provides the overall U-to-S delay for the initial Leiden cluster. Finally, the initial Leiden cluster with the highest U-to-S delay is considered the correct initiation. The DPT under this condition is used to initialize pseudotime for TSvelo. If multiple lineages are detected, TSvelo will model each lineage independently and subsequently merge them in the final step. A detailed explanation with example illustrating how lineages are segmented is provided in the **Fig. S10 in Supplementary Information**. Results of the initialization strategy on all datasets are shown in **Fig. S12**, demonstrating that clusters with the highest U-to-S delay scores can be reliably identified as the initial cell states. We also provide a simulation study to verify this initialization strategy in the **Fig. S2**.

### The ODE model in TSvelo

Suppose *u*_*g*_ and *s*_*g*_ are the abundance of unspliced and spliced RNA, and *α*_*g*_ (*t*), *β*_*g*_, *γ*_*g*_ are the transcription rate, splicing rate and degradation rate, respectively. The dynamics of a velocity gene g is modeled as:

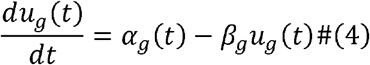

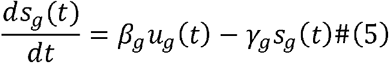

We assume that the gene amd cell-specific transcriptional rate *α*_*g*_ (*t*) is influence by the expression of Transcriptional Factors (TFs), while *β*_*g*_ and *γ*_*g*_ are gene-specific constant parameters. Considering the wide usage of linear models for are gene-relations in previous studies^27,34,37^ we *α*_*g*_ (*t*)

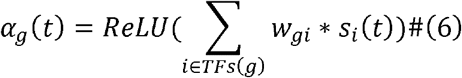

In order to include more TFs in the modeling, we reserve those TFs that are not selected as velocity genes. These TFs are excluded from the velocity genes selection because their unspliced-spliced expression does not provide sufficient information, primarily due to high noise in the unspliced RNA. Consequently, we incorporate these TFs, whose spliced abundance is denoted as

, into the TSvelo by directly modeling the process between transcriptional signal to mature RNA,

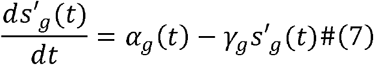

As a result, we can get the ODE model with parameters matrix *A*:

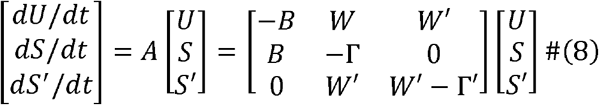

*U* = [*u*_1_,*u*_2_ …, *u*_*G*_]^*T*^ ∈ *R*^*G*^ = *S*^*’*^ *=* [*s*_G+1_, *s*_G+2_ …, *s*_G+*K*_]^*T*^ ∈ *R*^*K*^, *G* is the number of velocity genes and is the number of additional TFs which are not selected as velocity genes. *B* ∈ *R*^*G×G*^ and Γ ∈ *R*^*G×G*^ are the diagonal matrices consist of splicing rates [*β*_1_, *β*_*2*_, *… β*_*G*_] and degradation rates [*γ*_1_, *γ*_*2*_, *… γ*_*G*_] for velocity genes, respectively. Γ^’^ ∈ *R*^*K×K*^ is the diagonal matrices consist of degradation rates [*γ*_*G+*1_, *γ*_*G+2*_, *… γ*_*G+K*_] for these additional TFs. *W* ∈ *R*^*G×G*^ and *W’* ∈ *R*^*K×K*^ denote the matrices representing the TF-target relationships for the TFs selected as velocity genes and those not selected as velocity genes, respectively. Using these gene-gene weight matrices, TSvelo can model the gene- and cell-specific transcriptional rate *α*_*g*_. The whole parameters matrix *A* ∈ *R*^*(2G+K)ß (2G+K)*^

### Optimizing global time and Neural ODE in EM framework

In TSvelo, the dynamics described in **Eq. 7** are implemented using a neural ordinary differential equation (ODE) model. Given an initial state and the parameters in matrix *A*, the neural ODE model can compute the values of U and S at any time step. This approach eliminates the need for an analytical solution for unspliced and spliced expression, enhancing the flexibility of TSvelo. By default, the number of time steps in the ODE is set to 1,000. The ReLU activation function is applied to the parameters *α*_*g*_,*β*_*g*_, *γ*_*g*_, in order to prevent them from taking negative values. Notably, TSvelo does not incorporate deep neural networks or encoders. Since TSvelo models all genes simultaneously, we normalize the unspliced and spliced abundances of each gene by its standard deviation before inputting the data into the Neural ODE model to ensure that the influence of each gene is balanced.

TSvelo optimizes the pseudotime *t* and parameters matrix *A* in ODE iteratively using an Expectation-Maximization (EM) approach. The maximum number of iterations is set to 30 by default, with an early stopping criterion applied. Given the parameters in ODE system, we can compute the predicted values of unspliced (for velocity genes) and spliced (for velocity genes and additional TFs) at all time steps, denoted as 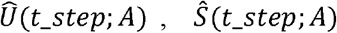 and 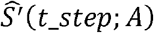, where *t*_*step*_ ∈{0,1,2…,999}. Given the time steps assignment *t*_*c*_ for each cell, we can get the expected values for velocity genes at all cells as 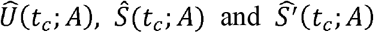. The loss function used in EM algorithm is the mean squared error between the velocity genes at all cells as and expected value with the observed data.

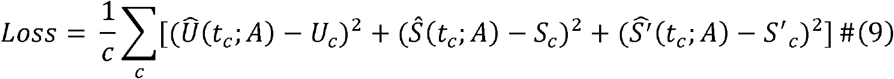

The EM algorithm will iteratively update the time assignment and parameters in neural ODE by minimizing the above loss function.

In the E-step, following the strategy adopted in scVelo model^6^, TSvelo updates the time step assigned to each cell using grid search with time steps range *t*_*step*_ ∈{0,1,2,…,999}, which is achieved by finding the minimum distance between the observed unspliced-spliced expression and the model prediction (*Û* and *Ŝ*) at all time steps.

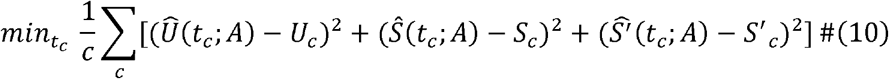

In the M-step, the parameter matrix *A* is updated using gradient descent within the neural ODE model to minimize the loss function. Here we use the prior gene regulation knowledge to constraint the weight matrix *W* and *W*^*’*^ in *A*. Only If gene *i* is annotated as a TF for gene *j* in the ENCODE or ChEA databases, the corresponding *w*_*ij*_ could be learnable. Otherwise is *w*_*ij*_ fixed at zero. The usage of prior knowledge could keep the matrix sparse and avoid overfitting. The neural ODE module is implemented using the torchdiffeq package and trained with the Adam optimizer, a learning rate of 0.05, and a maximum of 500 epochs, incorporating an early stopping criterion to prevent overfitting.

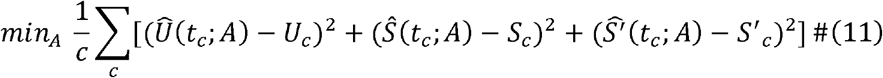

By performing the EM optimization, TSvelo could fit the high-dimensional unspliced-spliced data across multiple genes well, and learn the global pseudotime for each cell. After the training procedure, the RNA velocity could be calculated using those parameters in matrix *A* which could be further used for cell fate prediction.

### Metrics for evaluating

(1) Velocity consistency (VCon). We used the scvelo.velocity_confidence() function from scVelo to evaluate velocity consistency, interpreting the results as a measure of how consistent velocities are within neighboring cells. Velocity consistency is especially suitable for evaluating the RNA velocity modeling on single lineage. For each cell *c*, the velocity consistency is calculated as follows:

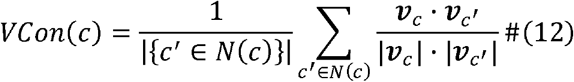

where *N*(*c*) represents the neighboring cells of a given cell *c*. *v*_*c*_ and 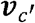 denote the low-dimensional velocity vectors of cell and its neighboring cell *c*^*’*^.

(2) Cross-boundary direction correctness (CBDir). Cross-boundary direction correctness is initially introduced by VeloAE^61^, which assesses the accuracy of transitions from a source cluster to a target cluster by examining the boundary cells, and requires ground truth annotations. We directly run the function unitvelo.evaluate() provided in UniTVelo^7^ to obtain the Cross-boundary direction correctness. In detail, the CBDir is calculated as follows:

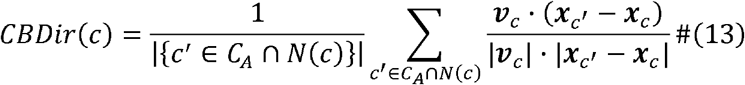

where *C*_*A*_ denotes the set of cells in the target cluster A, and *N*_(c)_ represents the neighboring cells of a given cell *c. v*_*c*_ and *x*_*c*_ denote the low-dimensional velocity and state vectors of cell *c*, respectively, and 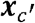 denotes the state vector of its neighboring cell.

(3) Within-cluster velocity coherence (ICCoh). Within-cluster velocity coherence is initially introduced by VeloAE^61^, which measures the coherence of velocities within a single cluster using a cosine similarity score between cell velocities. We applied the function unitvelo.evaluate() provided by UniTVelo^7^ to directly compute the within-cluster velocity coherence. Using the same notation as defined above, the CBDir is calculated as follows:

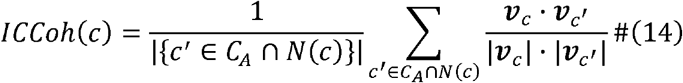

### The comparison between the 3D phase portrait and traditional 2D phase portrait

To quantitatively assess whether TSvelo can distinguish cell types, we evaluated the separability of cell-type labels in both the 2D (unspliced–spliced) phase portrait adopted by previous RNA velocity approaches, and the 3D (α–unspliced–spliced, *α*denotes the transcriptional rate) phase portrait introduced by TSvelo.

Specifically, we evaluated how well the embedding preserves cell-type information using a k-nearest neighbors (kNN) classification accuracy with 5-fold cross-validation. Given an embedding matrix in 2D or 3D space (*X* ∈ ℝ^*n*d*^, where *n* is the number of cells and is *d* is 2 or 3) and corresponding cell-type labels (*y* ∈ {1, …, *C*}), we partition the data into five folds. For each fold (*k*), a kNN classifier with *K*=5, denoted as *f*^(*k*)^, is trained on the training subset and evaluated on the held-out test subset. The classification accuracy for the *k*-th fold is defined as

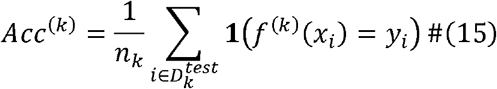

where *n*_*k*_ is the number of samples in the test set and **1** (·) is the indicator function. The final score is obtained by averaging across all folds:

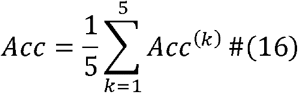

This metric directly assesses whether cells of the same type are positioned close to each other in the embedding space, and is widely used to quantify representation quality.

### GO term enrichment analysis

Gene Ontology (GO) terms enrichment analysis is a widely used method for identifying biological processes, molecular functions, or cellular components that are overrepresented in a given set of genes compared to a background set. Here we use the biological processes enrichment analysis in GO terms database to explore the associated biological roles of those selected velocity genes. To perform the GO term enrichment analysis, we utilized the “gseapy.enrichr()” function in Python with using Fisher’s exact test by default.

### Merging of lineage-specific results

For datasets containing multiple lineages, TSvelo applies its model independently to each lineage. The resulting lineage-specific outputs are integrated into a unified representation by aggregating both expression layers and cell-level annotations across branches.

All objects are aligned to a common gene space by restricting to the shared set of genes. Let *b* = 1, …, *B* index the lineage-specific results. For each cell *c* and each inferred quantity (e.g. velocity and psuedotime), TSvelo collects the corresponding results *x*_*c,b*_ from all lineages in which the cell is present. Layer values are merged on a per-cell basis using a weighted average, where the weight for each branch is given by its number of cells, *n*_*b*_. The merged value for cell *c* is computed as

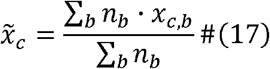

Continuous variables are averaged, while discrete variables are cast to integer values after aggregation.

Finally, the aggregated layer values and annotations are assigned back according to the global cell index, producing a unified dataset that integrates lineage-specific inferences.

## Data Availability Statement

The pancreatic endocrinogenesis dataset^62^ comprises the single-cell RNA-seq (10X) data of pancreatic epithelial and Ngn3-Venus fusion cells sampled from mouse embryonic day 15.5, which could be loaded using scVelo’s package scvelo.datasets.pancreas().

The gastrulation erythroid dataset, which is selected from the transcriptional profiles of mouse embryos^39^, which could be loaded using scVelo’s package scvelo.datasets.pancreas().

10× embryonic mouse brain dataset is provided at the 10x website at https://www.10xgenomics.com/resources/datasets/fresh-embryonic-e-18-mouse-brain-5-k-1-standard-1-0-0. The data preprocessed by Multivelo is utilized in this study, (https://multivelo.readthedocs.io/en/latest/MultiVelo_Fig2.html).

The dentate gyrus neurogenesis data is available at http://pklab.med.harvard.edu/velocyto/DentateGyrus/DentateGyrus.loom

The LARRY dataset has been shared by pyrovelocity, which could be accessed at https://figshare.com/articles/dataset/larry_invitro_adata_sub_raw_h5ad/20780344.

The raw data of Hindbrain (pons) of adolescent mice is from https://pklab.med.harvard.edu/ruslan/velocity/oligos/.

The ENCODE TF-target database^58^ website: https://maayanlab.cloud/Harmonizome/dataset/ENCODE+Transcription+Factor+Targets.

The ChEA TF-target database^59^ website: https://maayanlab.cloud/Harmonizome/dataset/CHEA+Transcription+Factor+Targets. The results of BayVel^11^ on the pancreas dataset is downloaded from its GitHub page at https://github.com/elenasabbioni/BayVel_notebooks/tree/main/real%20data/Pancreas/moments/output.

## Code Availability Statement

TSvelo is implemented in Python. The source code can be downloaded from the GitHub repository, https://github.com/lijc0804/TSvelo.

## Acknowledgement

This work was supported by the National Key R&D Program of China (2023YFF1204500 to Y.Y.), and the National Natural Science Foundation of China (No. 62503452 to J.L.).

