## Supplementary figures and tables for "TSvelo: Comprehensive RNA velocity by modeling the cascade of gene regulation, transcription and splicing"

**Table S1. The Comparison of RNA velocity approaches**

| **Methods** | **Biological scope** | **Key assumptions** | **Inference Framework** | **Modeling Granularity** |
| --- | --- | --- | --- | --- |
| velocyto | Transcription and splicing | Steady state. | Least squares regression | Per-gene |
| scVelo | Transcription and splicing | Two-step transcription rates. Constant splicing rates and degradation rates. | Expectation Maximization (EM) | Per-gene |
| VeloAE | Transcription and splicing | Steady state. | Auto-Encoder (AE) | Joint (all genes simultaneously) |
| UniTVelo | Transcription and splicing | Spliced RNA abundance is a twice-differentiable function of time | Radial Basis Function (RBF) | Per-gene |
| Dynamo | Transcription and splicing or metabolic | Two-step transcription rates. Constant splicing rates and degradation rates. | Ordinary Differential Equations (ODEs) and sparseVFC | Per-gene |
| cellDancer | Transcription and splicing | Cell-specific time-dependent transcription rates, splicing rates and degradation rates. | Convolutional Neural Network (CNN) | Per-gene |
| MultiVelo | Transcription and splicing | Two-step chromatin accessibility states. Constant splicing rates and degradation rates. | EM | Per-gene |
| BayVel | Transcription and splicing | A product of Poisson distributions for the initial state. | Markov Chain Monte Carlo | Per-gene |
| TFvelo | Regulation | Cell-specific time-dependent transcription rate. Constant degradation rates. | Generalized Expectation Maximization | Per-gene |
| TSvelo | Regulation, transcription and splicing | Cell-specific time-dependent transcription rates. Constant splicing rates and degradation rates. | Neural ODE | Joint (all genes simultaneously) |

**Simulation data analysis**
We generated synthetic single-cell RNA velocity datasets using a mechanistic transcriptional dynamics model with one or multiple developmental branches. The system included 200 genes, among which 30 were designated as transcription factors (TFs).

For each branch, we independently sampled a TF–target regulatory matrix $W\in\mathbb{R}^{30\times200}$ from a standard normal distribution to simulate distinct GRN structures. Gene expression dynamics were modeled using a coupled ordinary differential equation (ODE) system describing unspliced and spliced RNA abundances:

$$\frac{du_{g}\left( t \right)}{dt}=\alpha_{g}\left( t \right)-\beta_{g}u_{g}\left( t \right)$$

$$\frac{ds_{g}(t)}{dt}=\beta_{g}u_{g}\left( t \right)-\gamma_{g}s_{g}\left( t \right)$$

where $u$ and $s$ denote unspliced and spliced RNA levels, respectively. The transcription rate $\alpha$ was computed as a nonlinear function of TF expression, defined as a weighted sum of spliced TF abundance, followed by clipping to ensure bounded activation.

Each branch is initialized from the same randomly sampled initial condition drawn from a gamma distribution, allowing controlled divergence of trajectories driven solely by branch-specific regulatory programs.

To simulate observed sequencing counts, we introduced technical noise by scaling latent expression levels with cell-specific library sizes drawn from a log-normal distribution. The resulting expression counts were generated using a negative binomial sampling model:

$$X\sim NB(\mu=library size\times x,\theta)$$

where $\theta$ controls overdispersion, with smaller values corresponding to higher noise levels. The final datasets consist of paired unspliced (U) and spliced (S) count matrices with realistic transcriptional stochasticity and branching gene regulatory dynamics. For each branch, cells were further divided into three developmental stages for downstream analysis.

We evaluated TSvelo on multiple simulated datasets with varying numbers of branches and noise levels. There are two or three branches start from the same root cell groups in these datasets (Branch 1: stage 0 - stage 1 - stage 2. Branch 2: stage 0 - stage 3 - stage 4. Branch 3: stage 0 - stage 5 - stage 6). The results of initial state identification based on the unspliced-to-spliced (U-to-S) delay, along with the corresponding 2D velocity stream visualizations, are presented in **Fig. S1**. These results demonstrate that the U-to-S delay–based initialization is robust and consistently identifies cells corresponding to the earliest developmental stage (“stage 0”) across different simulation settings.

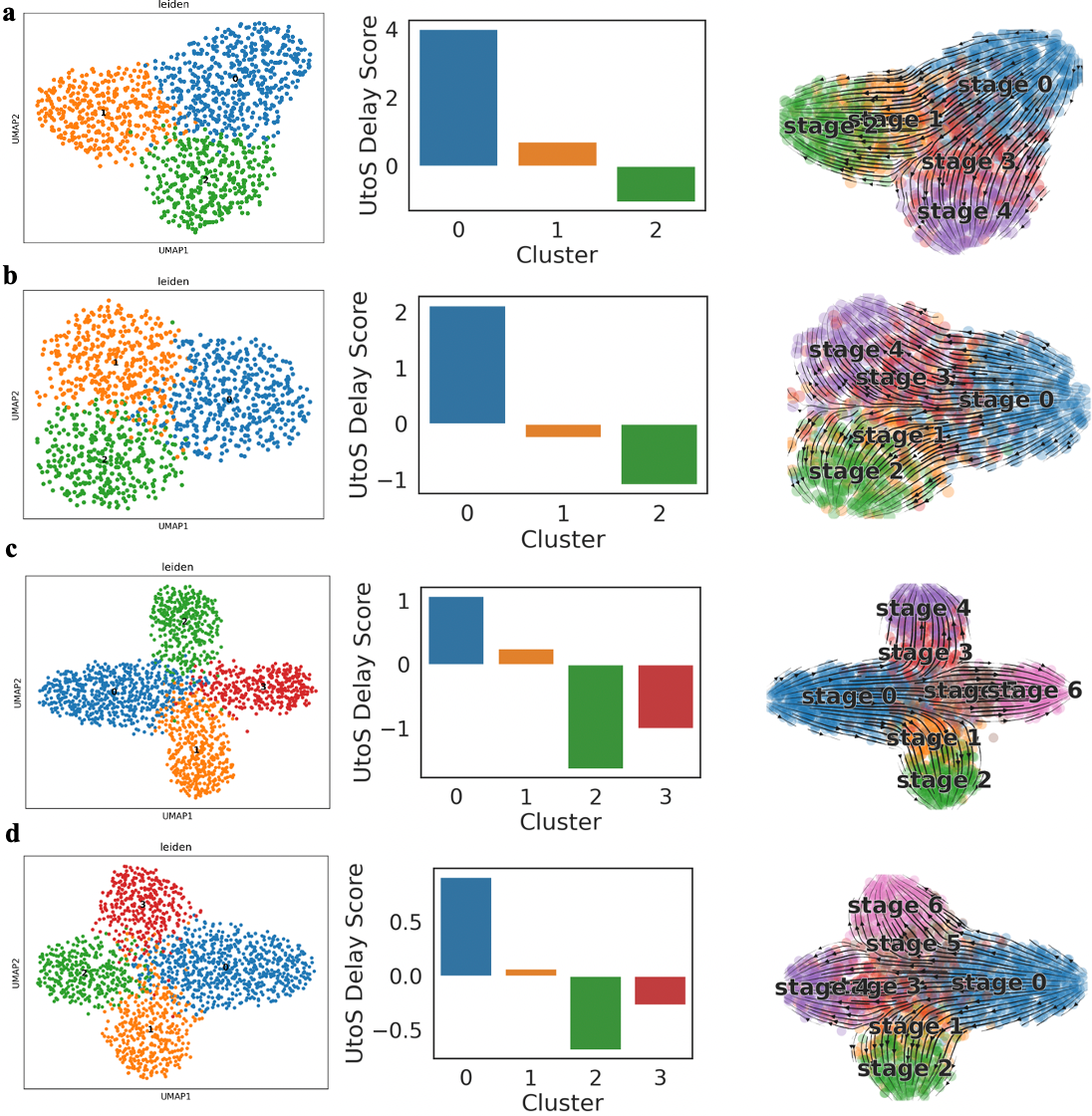

**Figure S1. Initial state detection and the 2D velocity stream obtained by TSvelo on multiple simulation datasets.**

We also evaluated TSvelo and those splicing-based RNA velocity approaches on multiple simulated datasets with varying numbers of branches and noise levels. There are one, two or three branches start from the same cell group in these datasets (Branch 1: stage 0 - stage 1 - stage 2. Branch 2: stage 0 - stage 3 - stage 4. Branch 3: stage 0 - stage 5 - stage 6). We primarily assessed performance using the cross-boundary direction correctness (CBDir) metric, as it directly evaluates inferred trajectories against ground-truth cell stage annotations. As shown in **Fig. S2**, TSvelo consistently achieves the highest accuracy across all simulation settings, particularly in scenarios with complex branching structures, which pose significant challenges for baseline methods.

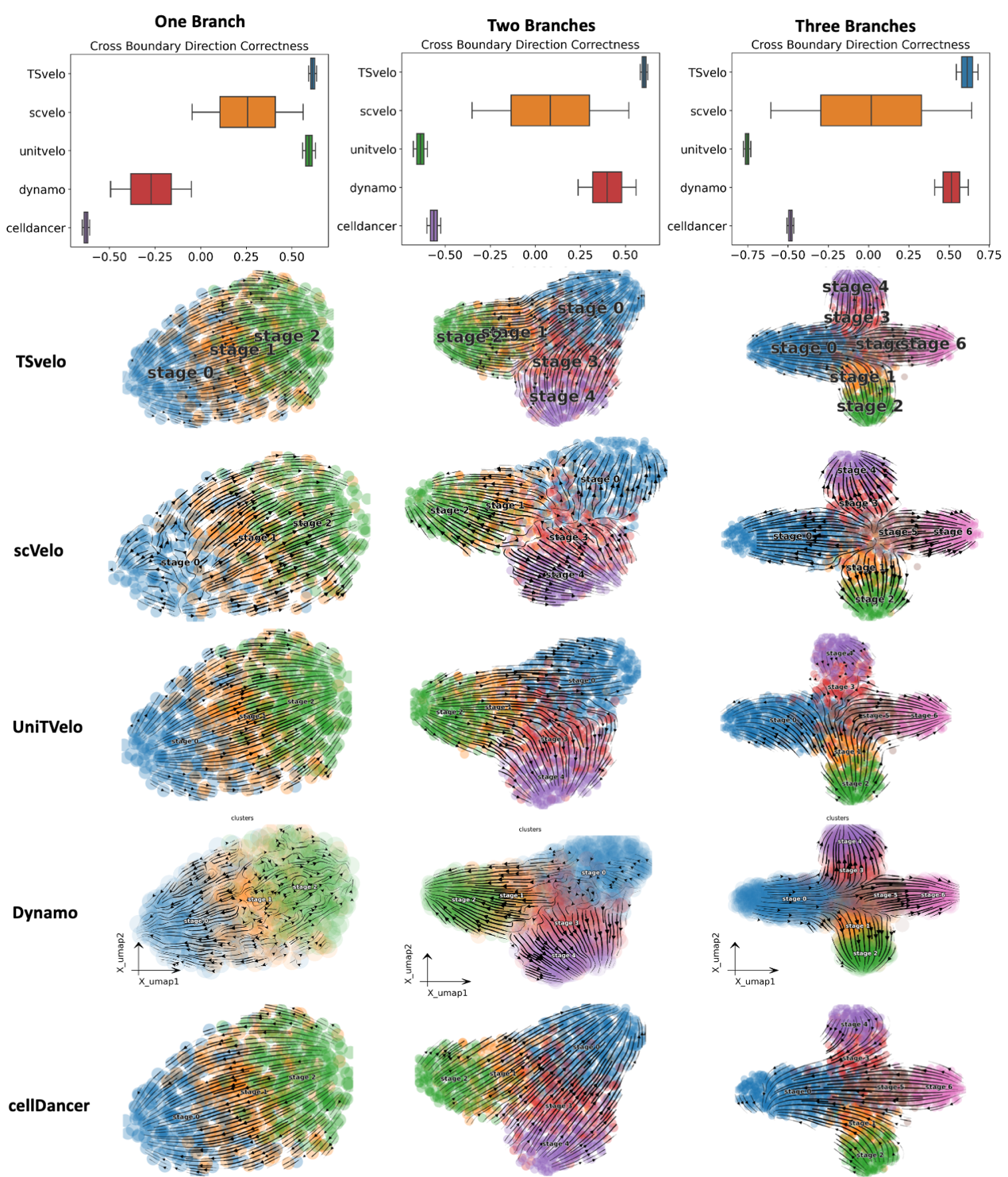

**Figure S2. Comparison between TSvelo and baseline approaches using the simulation data.**

**TSvelo can accurately model gene dynamics and cell fate on pons dataset**

The pons dataset originates from the hindbrain of adolescent mice and captures the differentiation pathway from oligodendrocyte precursor cells (OPCs) to committed oligodendrocyte precursor cells (COPs), progressing to newly formed oligodendrocytes (NFOLs), and finally culminating in myelin-forming oligodendrocytes (MFOLs). The velocity genes selected through TSvelo’s preprocessing strategies are mostly enriched in GO terms related to the development of nervous system, including regulation of axon extension involved in axon guidance (GO:0048841), axonogenesis (GO:0007409) and nervous system development (GO:0007399) (**Fig. S3a**).

Using the unspliced-to-spliced delay, TSvelo can correctly detect the initial Leiden cluster on pons dataset (**Fig. S3b** and **Fig. S3c**). After performing the model optimization for the neural ODE system, the pseudotime and velocity learned by TSvelo can accurately fit the differentiation process in the data (**Fig. S3d** and **Fig. S3e**). **Fig. S3f** illustrates the dynamics of multiple genes modeled by TSvelo. A challenge in this dataset is the abrupt shift in gene expression between OPCs (red) and COPs (orange). For example, as shown in **Fig. S3f** for CHN2 and DPYSL2, there is a significant gap in expression levels between OPCs and COPs. Despite this, TSvelo successfully captures their dynamic patterns.

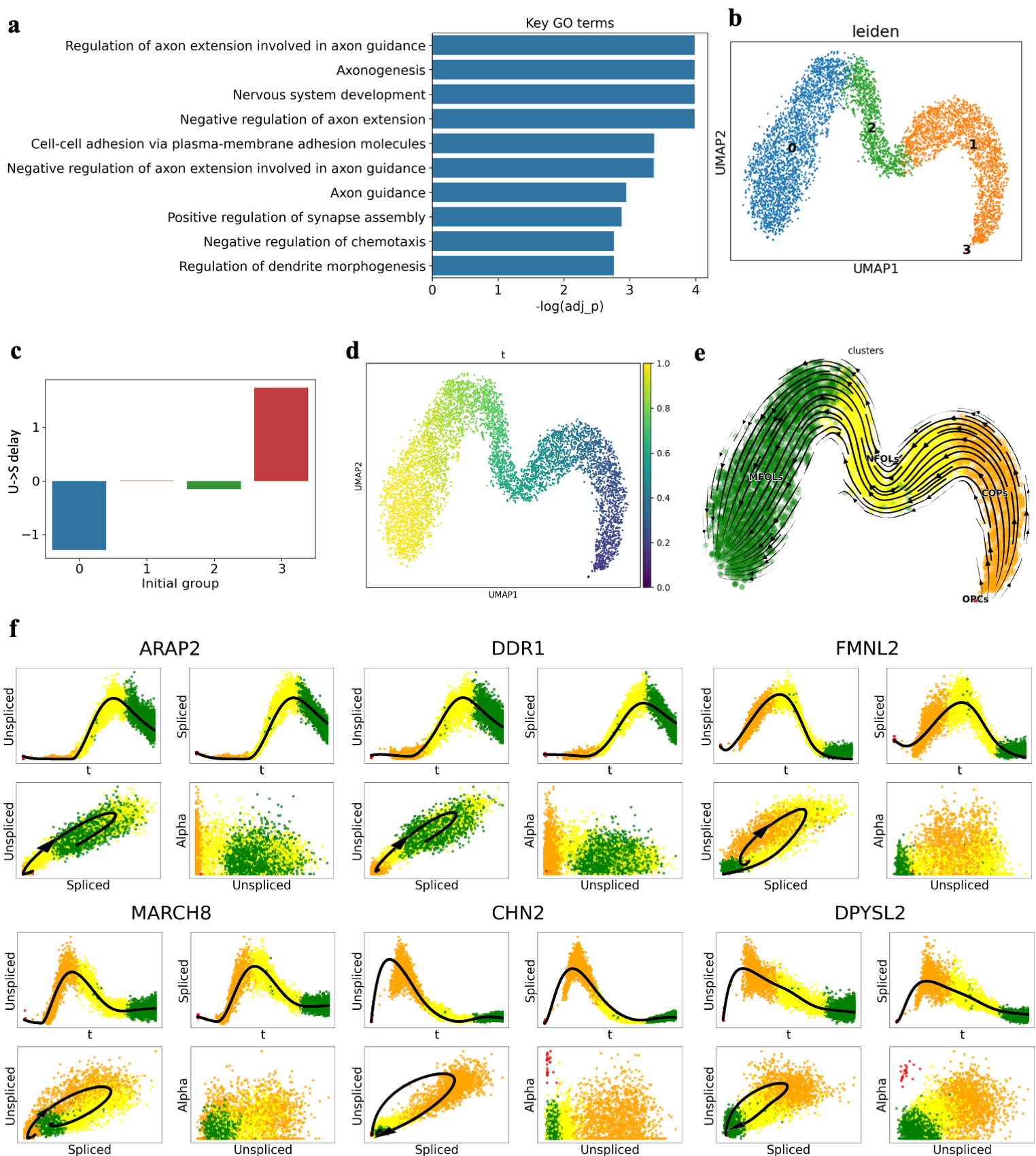

**Figure S3. Results on pons dataset.** (**a**) The GO terms which are mostly enriched in the selected velocity genes of TSvelo. (**b**) The Leiden clustering. (**c**) The U-to-S delay under different initial cluster choices. (**d**) The pseudotime learned with TSvelo. (**e**) The Stream-plot for visualization the RNA velocity inferred by TSvelo. (**f**) Dynamics fitting for multiple genes on the pons dataset. For each gene, four plots are displayed in a 2x2 layout: the u-t, s-t, u-s, and alpha-u plots.

**Additional results on pancreas dataset**

we conducted comparisons across all methods on the pancreas dataset, with quantitative evaluations shown in **Fig. S4**. In each plot, methods are ranked in descending order of their mean values. Numbers at the bottom indicate the sample size for each metric. Statistical significance is assessed using a one-sided Mann–Whitney U test, where *****, ***, **, and * denote p < 0.00001, 0.0001 ≤ p < 0.001, 0.001 ≤ p < 0.01, and 0.01 ≤ p < 0.05, respectively.

TSvelo significantly outperforms all baseline methods in terms of velocity consistency. For in-cluster coherence, TSvelo performs comparably to the best-performing baselines (UniTVelo and TFvelo) and significantly outperforms several competing methods, including CellDancer, Dynamo, and scVelo. For cross-boundary direction correctness, TSvelo shows consistent improvements in mean performance. The pairwise comparisons on cross-boundary direction correctness do not reach statistical significance. It is likely influenced by the limited number of independent samples (n = 7), which reduces statistical power for detecting differences. Importantly, TSvelo still achieves the best average performance among all methods, indicating a consistent overall trend in favor of TSvelo.

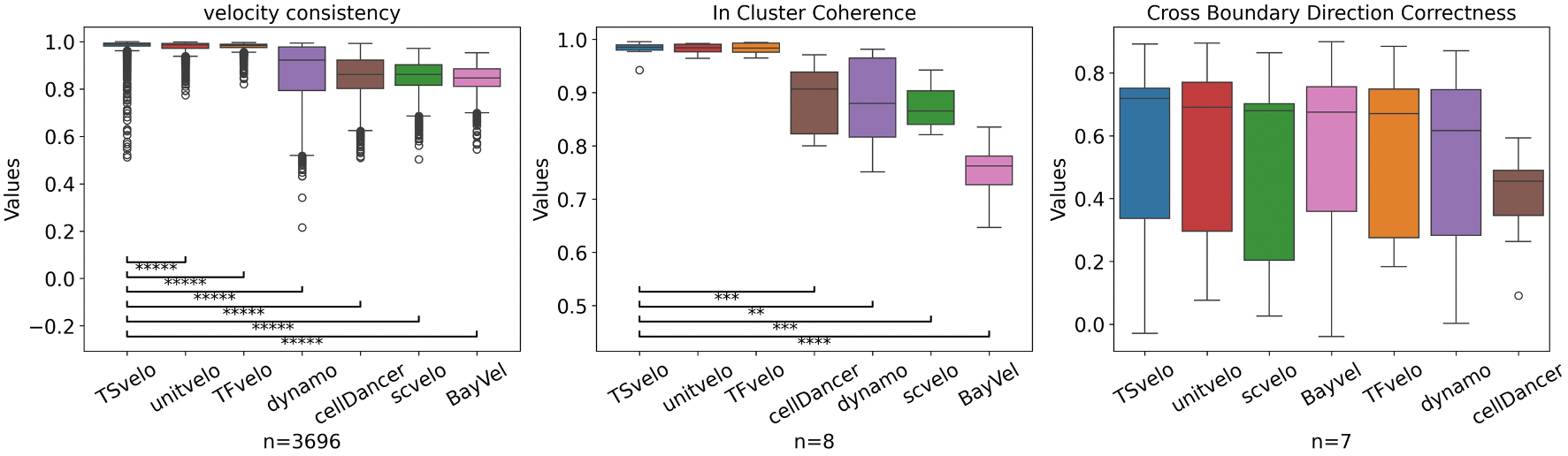

**Figure S4. The velocity consistency, in-cluster coherence and cross boundary direction correctness on pancreas dataset.** In each plot, methods are ranked in descending order of their mean values. Numbers at the bottom indicate the sample size for each metric. Significance is determined using a one-sided Mann–Whitney U test. *****, ****,***, ** and * represent p < 0.00001, 0.00001 ≤ p < 0.0001, 0.0001 ≤ p < 0.001, 0.001 ≤ p < 0.01, and 0.01 ≤ p < 0.05, respectively.

**Fig. S5** shows the dynamics fitting with TSvelo of more velocity genes on the pancreas dataset.

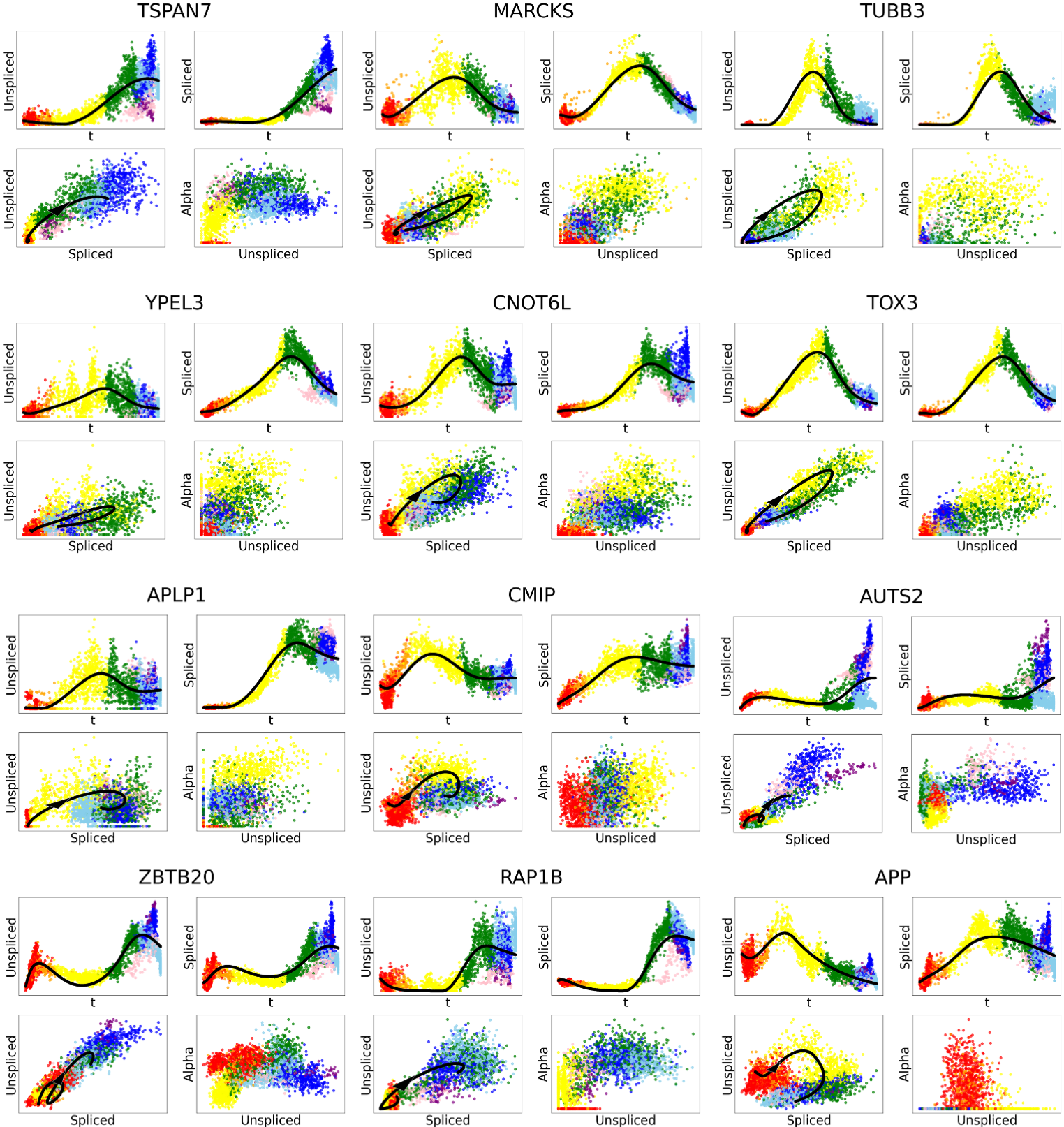

**Figure S5. Dynamics fitting of velocity genes on the pancreas dataset.** For each gene, four plots are displayed in a 2x2 layout: the u-t, s-t, u-s, and alpha-u plots.

TSvelo is capable of accurately modeling the dynamics of spliced mRNA abundance for those TFs which are not selected as velocity genes during preprocessing. These TFs cannot be well modeled in the unspliced–spliced phase portrait. However, TSvelo overcomes this limitation by directly modeling the process from transcription to spliced mRNA. **Fig. S6** shows the modeling on those TFs on pancreas dataset, where TSvelo directly models the dynamic between transcription and spliced abundance.

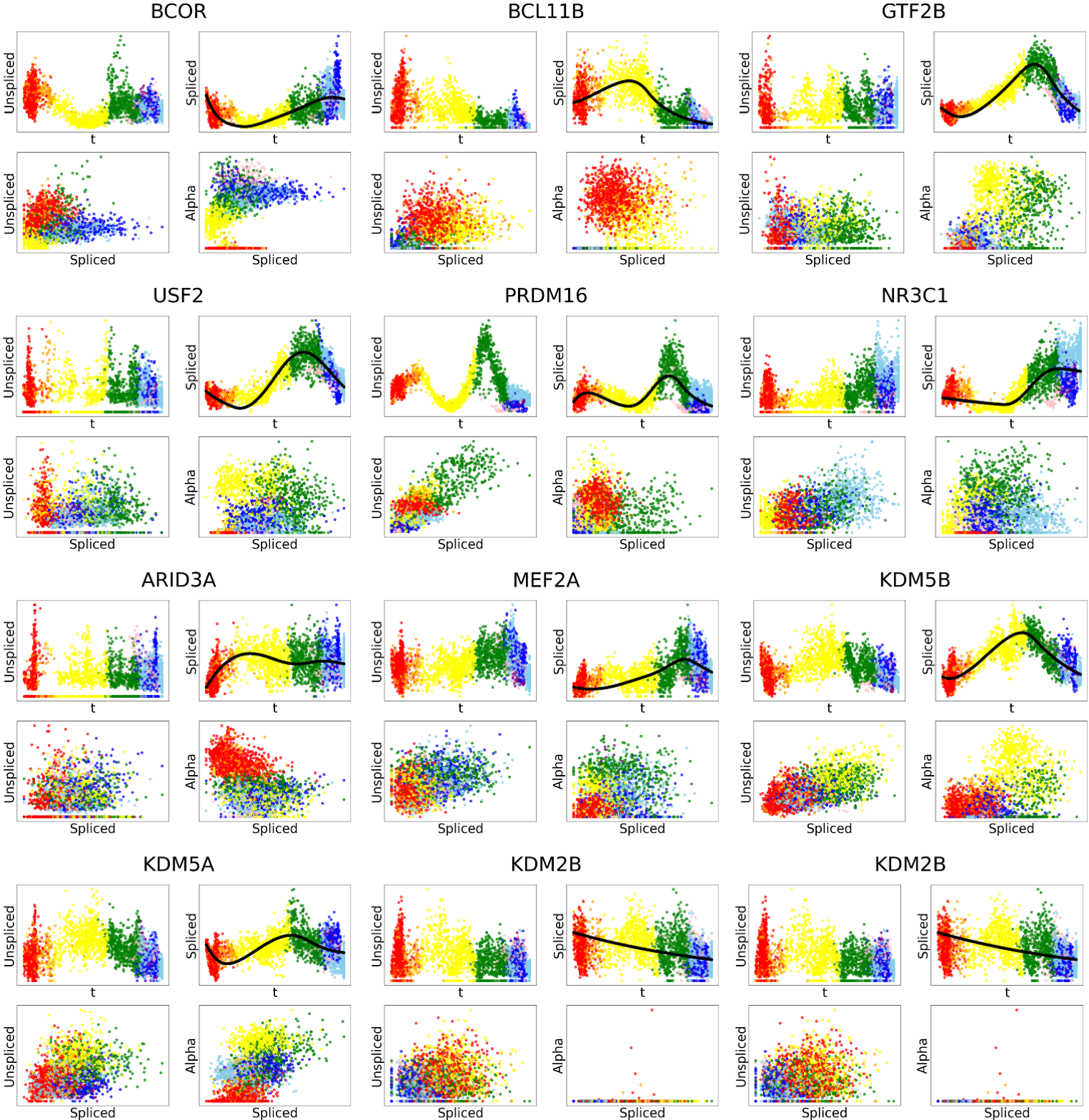

**Figure S6. Dynamics fitting for additional TFs on the pancreas dataset.** TSvelo directly models the dynamic between transcription and spliced abundance on these genes. For each gene, four plots are displayed in a 2x2 layout: the u-t, s-t, u-s, and alpha-u plots.

**Additional results on gastrulation erythroid dataset**

In the conventional 2D u–s phase portrait, cells from different transcriptional states may overlap, leading to reduced separability. In contrast, introducing the latent variable α expands the representation to a 3D space, which helps disentangle these mixed states and yields a clearer phase structure. We provide quantitative evidence on this gastrulation erythroid dataset in **Fig. S7**, showing that the 3D representation achieves consistently higher kNN classification accuracy for cell state separation compared to the 2D u–s embedding (one-sided Mann–Whitney U test, p-value = 0.002).

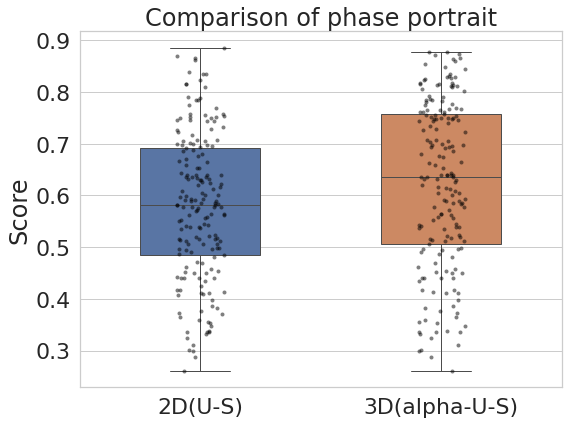

**Figure S7. The evaluation of the separability of cell-type labels in both the 2D (unspliced–spliced) phase portrait and the 3D (α–unspliced–spliced) phase portrait for the gastrulation erythroid dataset.**

**Fig. S7** shows the dynamics fitting on additional genes with TSvelo on the gastrulation erythroid dataset.

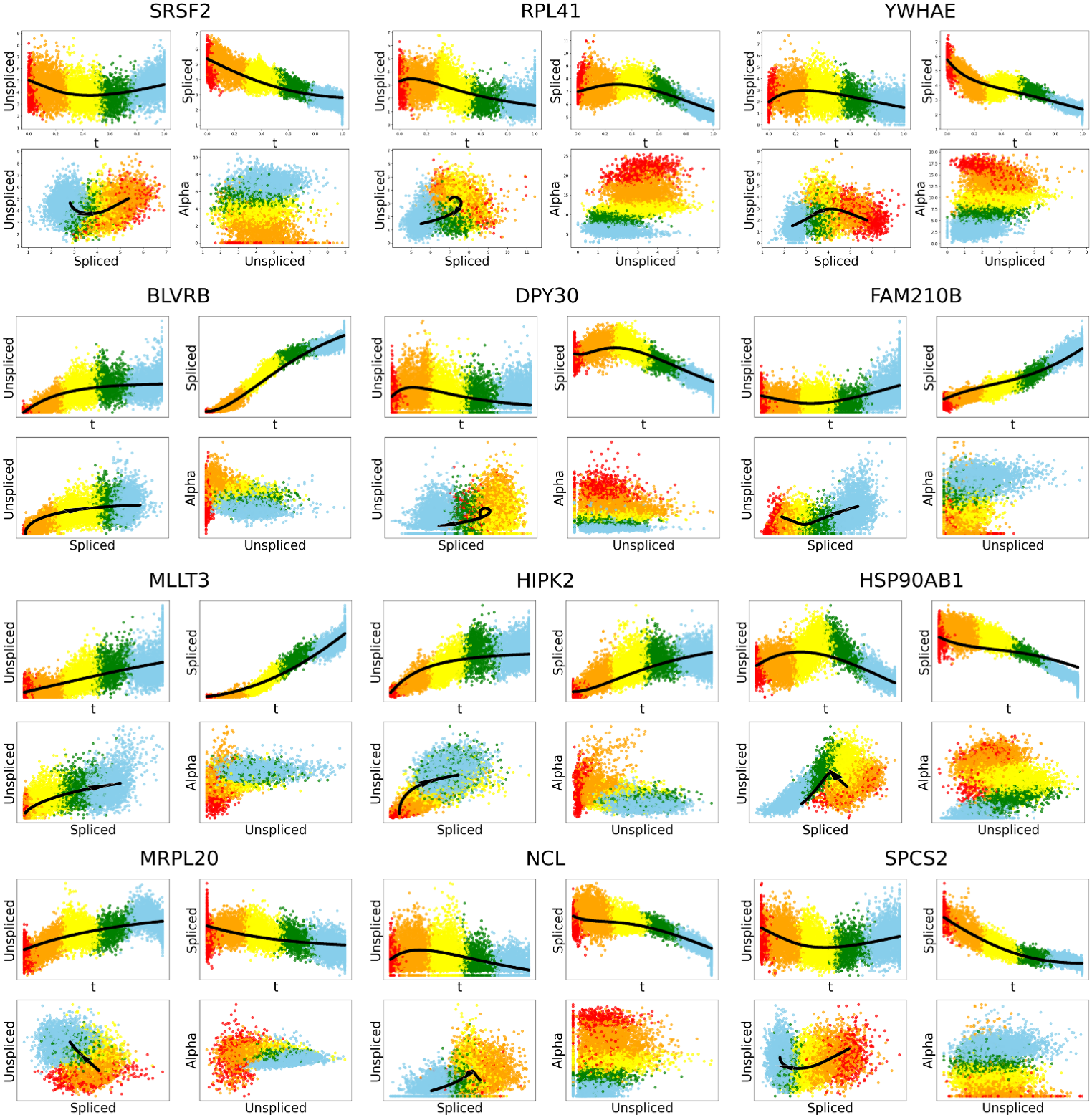

**Figure S8. Dynamics fitting of velocity genes on the gastrulation erythroid dataset.** For each gene, four plots are displayed in a 2x2 layout: the u-t, s-t, u-s, and alpha-u plots.

**Additional results on mouse brain dataset**

**Fig. S9** shows the dynamics fitting on additional genes with TSvelo on the mouse brain dataset.

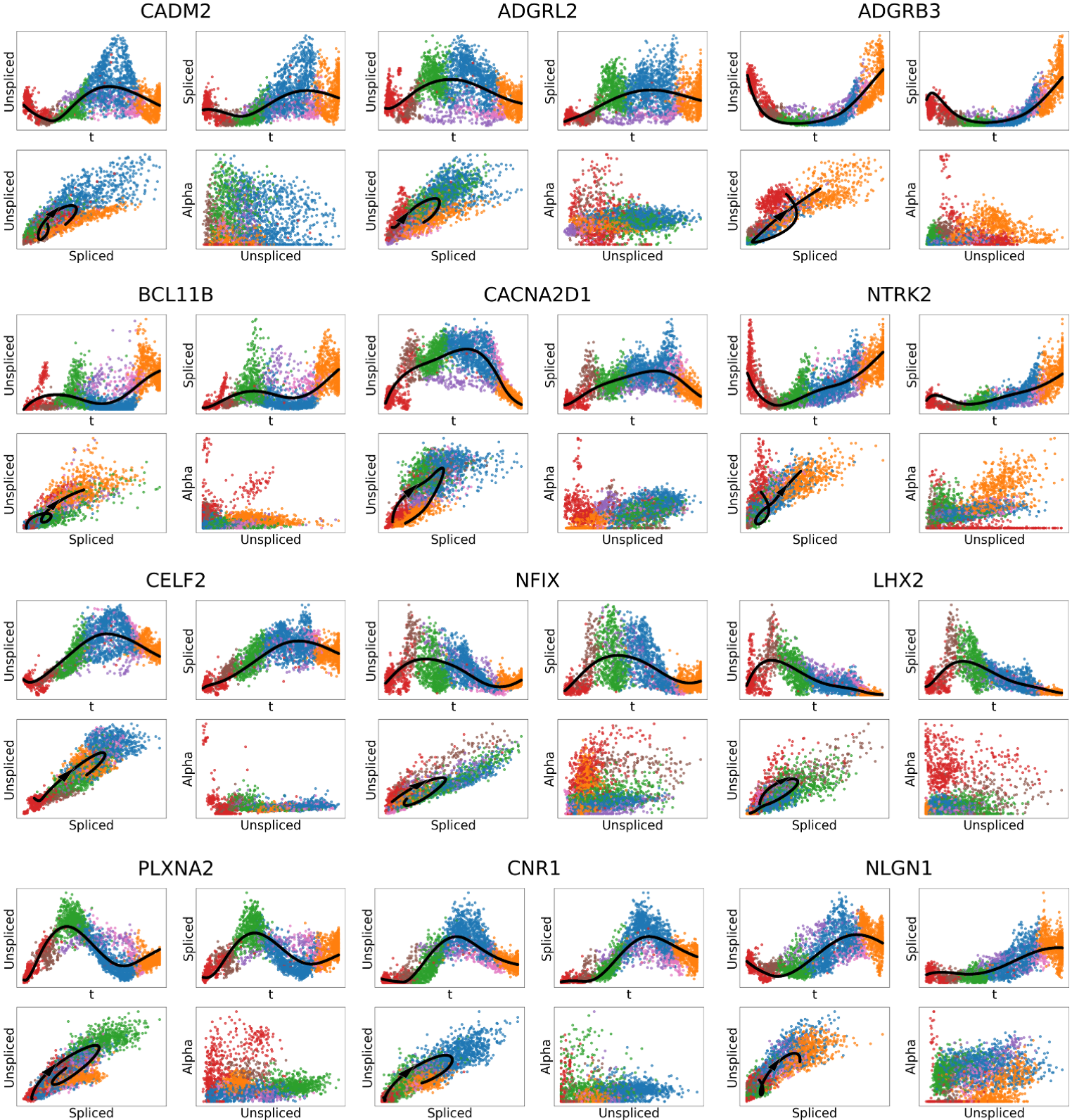

**Figure S9. Dynamics fitting for additional genes on the mouse brain dataset.** For each gene, four plots are displayed in a 2x2 layout: the u-t, s-t, u-s, and alpha-u plots.

**The lineages segmentation and combination on dentate gyrus dataset**

Using the dentate gyrus dataset, we demonstrate how lineages are segmented during preprocessing, and subsequently combined during postprocessing after being modeled individually. Notably, the lineage segmentation process is fully automated and does not require prior knowledge about the presence of multiple branches.

First, Leiden clustering is performed on the preprocessed scRNA-seq data, with the resolution set to 0.1 by default (**Fig. S10a**). A cluster-cluster graph is then constructed using PAGA (**Fig. S10b**), followed by a filtering strategy that removes edges with weights below a threshold of 0.02 by default. Next, each Leiden cluster is treated as an initial cluster, and the appropriate initialization is determined using the unspliced to spliced delay.

For instance, using Leiden cluster 3 (colored in red at **Fig. S10a**) as the initial cluster, diffusion pseudotime (DPT) is initialized with the selected cluster (**Fig. S10c**). Starting from this initial cluster, the shortest paths to all other clusters are computed. Any path that is a subset of another is discarded. The remaining paths correspond to the detected lineages. In the dentate gyrus dataset, initializing with Leiden cluster 3 yields three distinct lineages (**Fig. S10d**). By applying DPT to each lineage, we can calculate the U-to-S delay for each lineage and determine the overall U-to-S delay for this initialization with the selected cluster.

After calculating the U-to-S delay for each Leiden cluster initialization, the best initialization cluster is chosen based on the highest U-to-S delay (**Fig. S10e**). The corresponding DPT is then used to initialize pseudotime in the downstream neural ODE model. Subsequently, each lineage is processed independently in the TSvelo for optimization, yielding both the optimized pseudotime and velocity model. The results for each lineage are shown in **Fig. S10f**.

Finally, on the dataset where multiple branches are detected, to combine the models for all lineages and obtain the overall pseudotime and velocity stream, we compute the average velocity (**Fig. S10g**) and pseudotime (**Fig. S10h**) across the different lineages. For example, in the dentate gyrus dataset, since Leiden clusters 0, 1, 2, and 4 each belong to a single lineage, the velocity and pseudotime values calculated for these lineages are directly assigned to the combined file. For Leiden cluster 3, which is included in all three branches, the mean velocity and mean pseudotime across the three lineages are calculated for cells in this cluster.

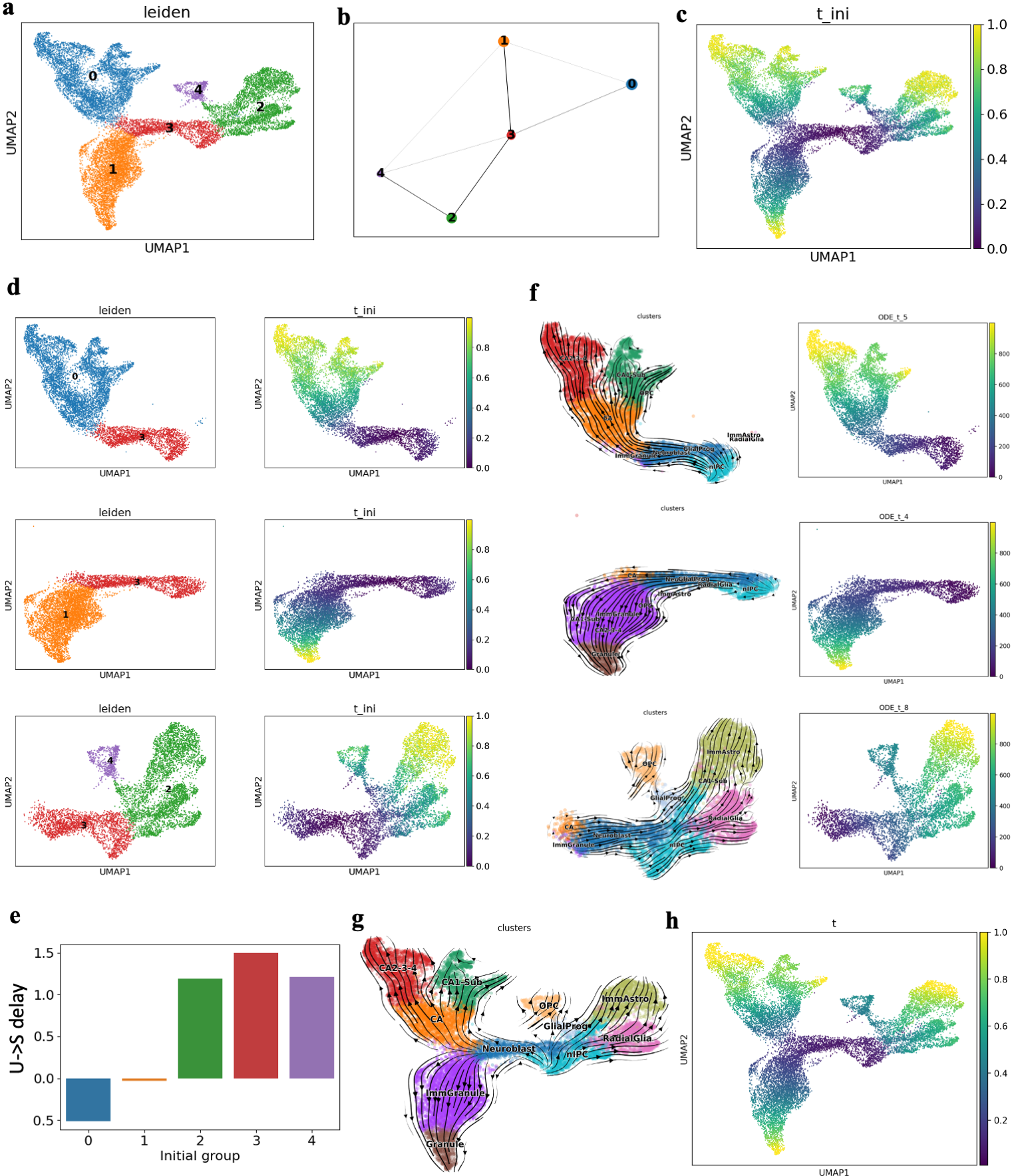

**Figure S10. Lineages segmentation on dentate gyrus dataset.** (**a**) The Leiden clustering. (**b**) The cluster-cluster graph constructed with PAGA. (**c**) The diffusion pseudotime when choosing initial cluster 3 as the initial one. (**d**) The three branches detected when choosing initial cluster 3 as the initial one. (**e**) U-to-S delay under different initial cluster choices. (**f**) The velocity stream and pseudotime inferred by TSvelo on each lineage. (**g**) The velocity stream obtained with the combined velocities from each lineage. (**h**) The combined TSvelo pseudotime from each lineage.

**Additional results on dentate gyrus dataset**

**Fig. S11** shows the detailed dynamics fitting on each lineage for ANK3, MAP1B and SLC1A2 on the dentate gyrus dataset.

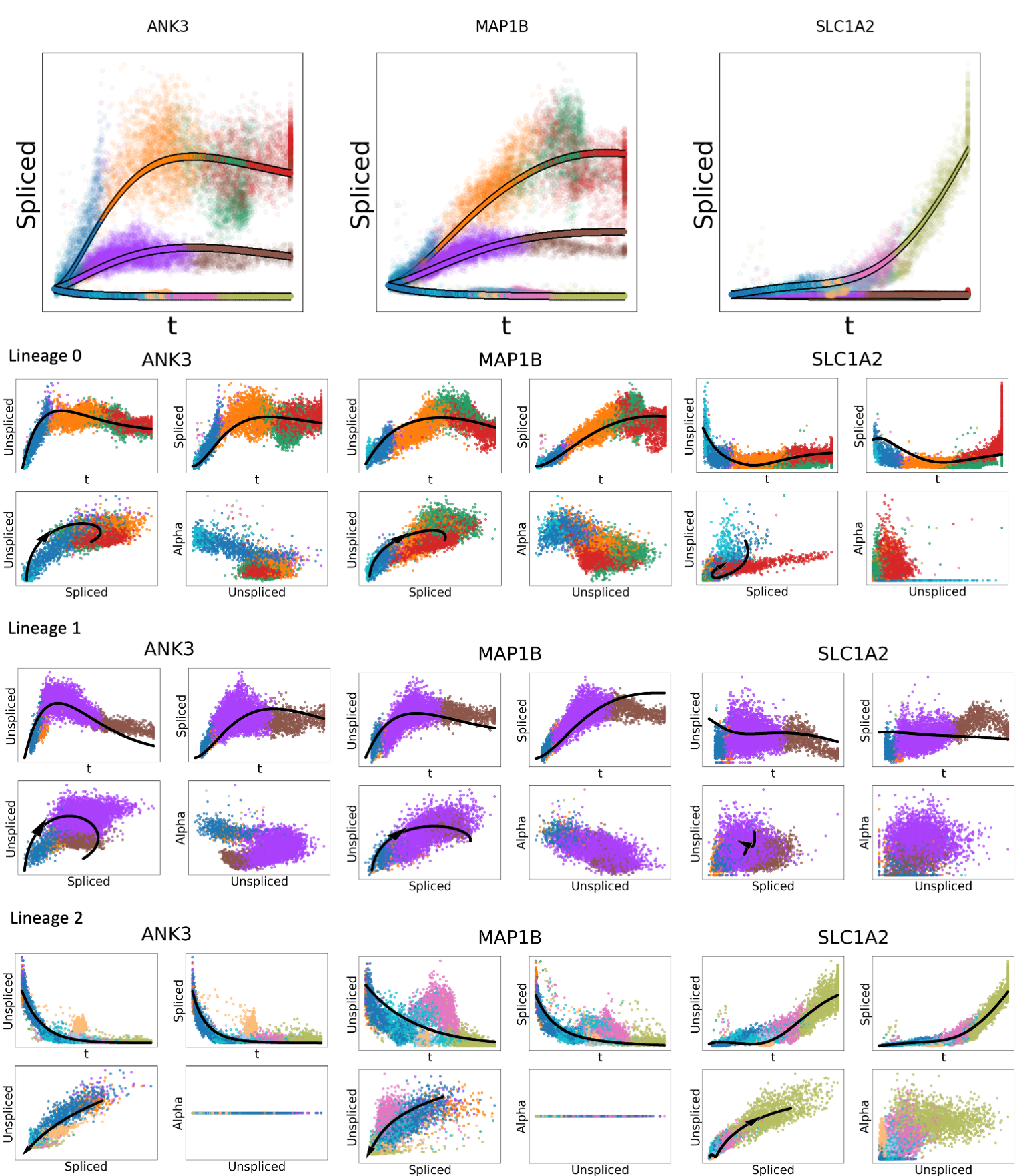

**Figure S11. Dynamics fitting for genes ANK3, MAP1B and SLC1A2 on each lineage of the dentate gyrus dataset.** For each gene, four plots are displayed in a 2x2 layout: the u-t, s-t, u-s, and alpha-u plots.

**The U-to-S delay scores for each Leiden cluster when treated as the initial state across all datasets.**

Along a correctly oriented trajectory, unspliced (U) expression is expected to precede spliced (S) expression due to transcriptional dynamics. Ideally, this U-to-S delay would be observable at the level of individual genes. However, due to the high noise inherent in scRNA-seq data, such delays are often not consistently detectable on a per-gene basis. To address this, we aggregate U-to-S delay signals across all genes and determine the lineage orientation by maximizing a global delay score. Under this criterion, the cluster from which all outgoing lineages exhibit the highest aggregated U-to-S delay is inferred to correspond to the initial state. The results on all datasets (**Fig. S12**) suggest that the highest U-to-S delay scores can be used to detect the initial cluster.

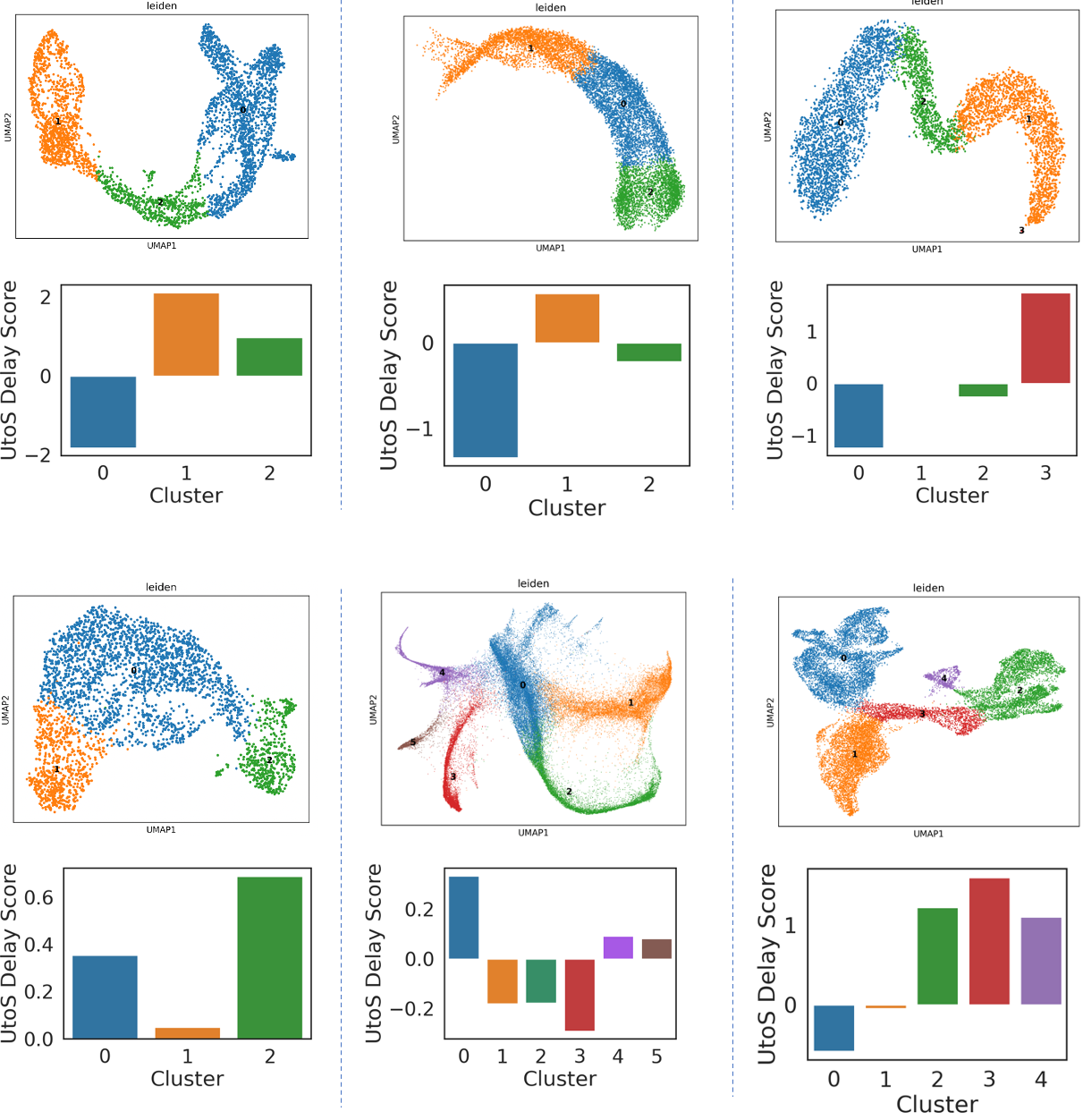

**Figure S12. The U-to-S delay scores for each Leiden cluster when treated as the initial state across all datasets.**

**Ablation studies on the selection of TF–target resources**

we additionally incorporated the DoRothEA regulon database as an alternative prior with confidence-level filtering. We further performed ablation studies on the pancreas dataset and the gastrulation erythroid dataset using different TF–target resources, including ChEA, ENCODE, and their combinations with DoRothEA.

The results on the pancreas dataset and the gastrulation erythroid dataset are shown in **Fig. S13** and **Fig. S14** respectively, which come up with the same conclusion. We observed highly consistent results across most TF–target prior combinations, including ChEA, ENCODE, ChEA+ENCODE, ChEA+DoRothEA, ENCODE+DoRothEA, and ChEA+ENCODE+DoRothEA. Using the pancreas dataset as example, the mean velocity consistency ranged from 0.985 to 0.995, the mean in-cluster coherence ranged from 0.983 to 0.992, and the mean cross-boundary direction correctness ranged from 0.719 to 0.740 across all settings. These consistently high and tightly bounded metrics indicate that TSvelo is largely insensitive to the specific choice of TF–target prior.

The only configuration showing reduced stability was the use of DoRothEA alone, particularly in terms of cross-boundary direction correctness. This is likely due to its comparatively limited coverage of TF–target interactions. For instance, in the pancreas dataset, only 81 out of 2000 highly variable genes (HVGs) could be associated with TFs based on DoRothEA, corresponding to 102 TF–target links in total, which may restrict downstream regulatory modeling. In contrast, ChEA covered 1793 genes with 13,976 TF–target links, and ENCODE covered 1854 genes with 33,076 links.

These results further suggest that integrating multiple TF–target resources can improve performance by increasing regulatory coverage and reducing false negatives in the prior network. Given that unsupported edges can be down-weighted during training, TSvelo appears to be more sensitive to missing true regulatory interactions than to the inclusion of spurious ones.

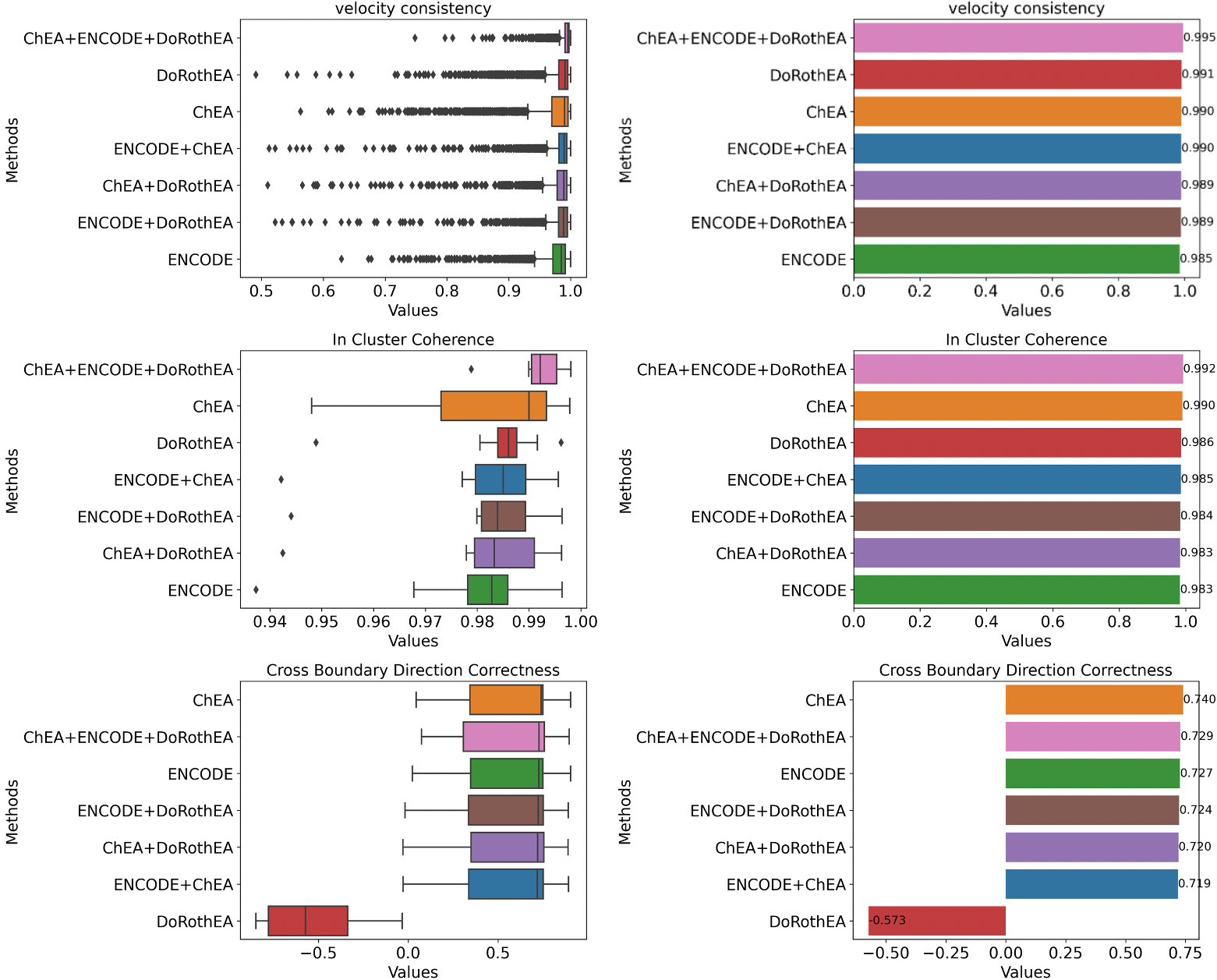

**Figure S13. The quantitative comparison of TSvelo modeling results on the pancreas dataset using different TF–target prior database.** The panels in the left column show the boxplot of velocity consistency, the in-cluster coherence and the cross-boundary direction correctness, respectively. The panels in the right column show the barplot of their mean values.

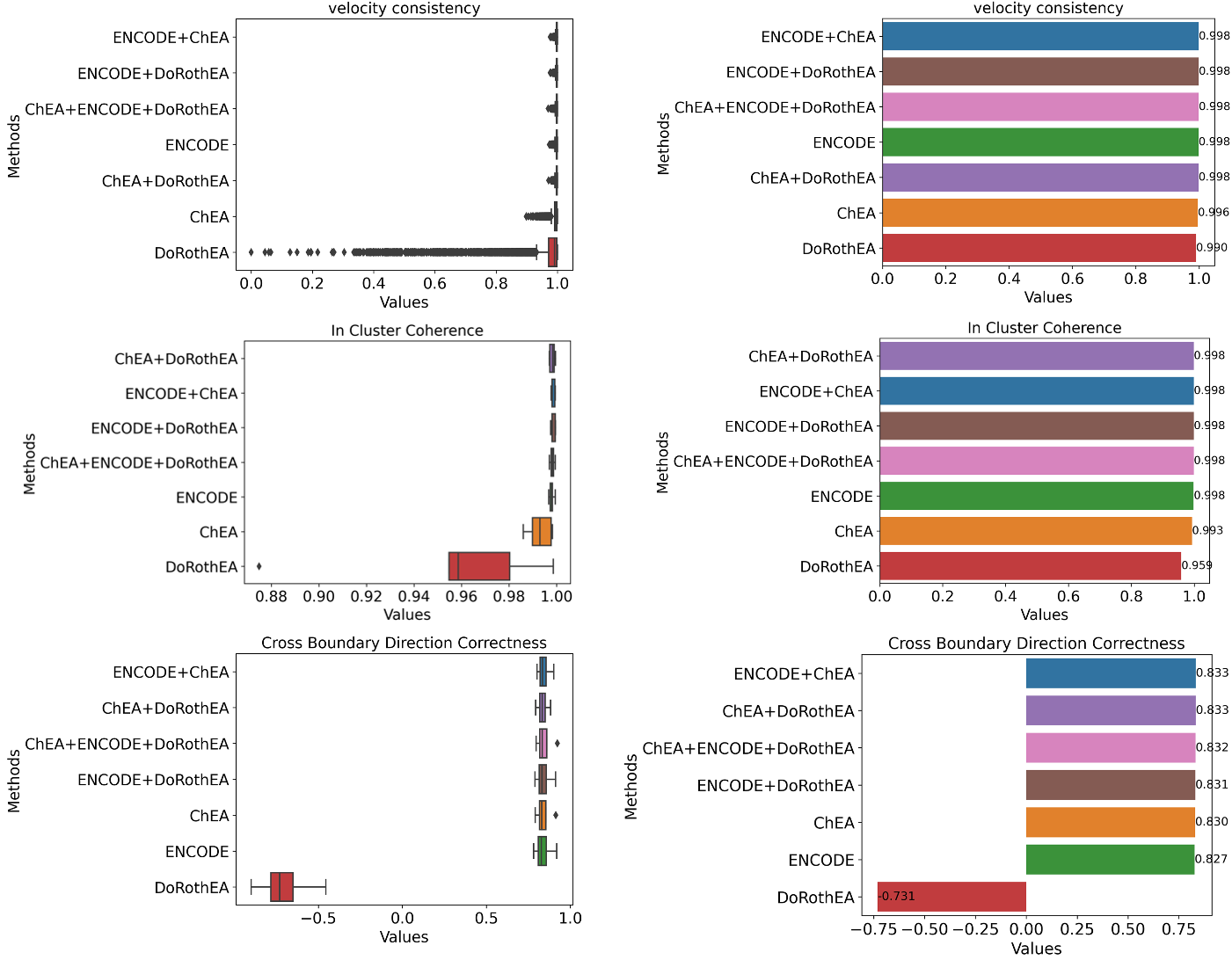

**Figure S14. The quantitative comparison of TSvelo modeling results on the gastrulation erythroid dataset using different TF–target prior database.** The panels in the left column show the boxplot of velocity consistency, the in-cluster coherence and the cross-boundary direction correctness, respectively. The panels in the right column show the barplot of their mean values.

**Comparison between the chrome accessibility rate used in MultiVelo and the learned transcription rate from TSvelo**

We have conducted the requested analysis by computing gene-wise chrome accessibility rate used in MultiVelo and the learned transcription rate from TSvelo, and evaluated their correlation across genes. As shown in **Fig. S15**, the two estimates exhibit almost no global correlation across genes, indicating that they capture substantially different aspects of regulatory information.

This discrepancy is not unexpected and reflects the fundamental differences between these modalities. scATAC-seq measures chromatin accessibility, which provides a proxy for cis-regulatory potential of genomic regions. However, ATAC signals are inherently sparse and often exhibit a near-binary structure, limiting their ability to directly capture fine-grained temporal regulatory dynamics. In contrast, TF RNA expression reflects downstream transcriptional output, which is shaped by multiple regulatory layers, including post-transcriptional regulation, protein activity, temporal delays, and indirect regulation through intermediate transcriptional or signaling pathways. As a result, these two modalities are expected to capture complementary but not directly comparable aspects of gene regulation.

Overall, this result suggests that ATAC-based and TF RNA-based signals capture distinct aspects of gene regulation. This further implies that integrating both modalities may be beneficial for future models that aim to more comprehensively characterize transcriptional regulation. We have added this discussion to the supplementary information.

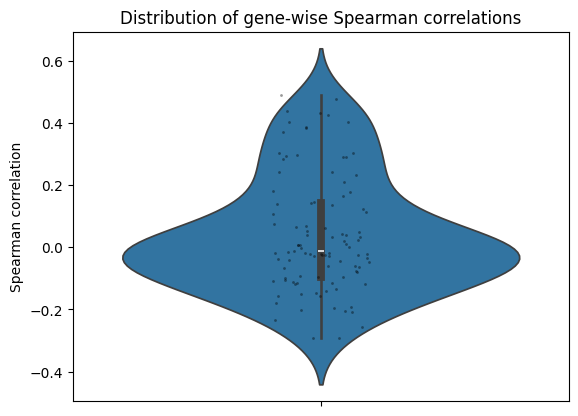

**Figure S15. The distribution of gene-wise Spearman correlation between the estimated transcription rate in TSvelo and the chrome accessibility rate in Multivelo.**

**Computational benchmark**

we have added a systematic comparison of runtime and GPU memory usage across TSvelo and ba methods using simulated datasets of increasing scale (600, 1200, and 1800 cells) on our NVIDIA GeForce RTX 3090 device with 24 GB memory.

**Table R2** shows differences in computational efficiency and resource requirements among methods. Specifically, classical methods such as scVelo and Dynamo exhibit very fast runtimes (10–24 seconds) and do not rely on GPU acceleration, reflecting their relatively lightweight modeling strategies. In contrast, deep learning–based approaches, including UniTVelo, cellDancer, and TSvelo, have higher computational costs due to their increased model complexity.

TSvelo exhibits a stable GPU memory footprint (~1.26 GB) across different dataset sizes, indicating that its memory usage is primarily determined by model architecture rather than the number of cells. This level of memory consumption is well within the capacity of modern GPUs and does not pose practical limitations. In terms of runtime, TSvelo scales approximately linearly with dataset size. The higher computational cost of TSvelo is mainly due to its EM-style optimization procedure, where each M-step also involves multiple optimization updates to infer gene regulatory effects in a global model. This design enables TSvelo to explicitly incorporate regulatory priors and jointly model gene interactions, which is not supported by these baseline methods.

**Table R2. Comparison of memory and runtime cost of different methods.**

|  | **Dataset scale** | **600 cells** | **1200 cells** | **1800 cells** |
| --- | --- | --- | --- | --- |
| **scVelo** | GPU Memory | - | - | - |
|  | Modeling Time | 10 sec | 12 sec | 14 sec |
| **Dynamo** | GPU Memory | - | - | - |
|  | Modeling Time | 24 sec | 20 sec | 24 sec |
| **cellDancer** | GPU Memory | - | - | - |
|  | Modeling Time | 82 sec | 118 sec | 245 sec |
| **UniTVelo** | GPU Memory | 930 MB | 994 MB | 1122 MB |
|  | Modeling Time | 345 sec | 637 sec | 1016 sec |
| **TSvelo** | GPU Memory | 1262 MB | 1264 MB | 1264 MB |
|  | Modeling Time | 734 sec | 1544 sec | 3027 sec |

To further improve runtime efficiency, TSvelo allows flexible control of the number of EM iterations. As shown in **Fig. S16** and **Table S3**, we evaluated performance under different iteration settings on the simulation dataset. The early stopping strategy employed in the EM framework of TSvelo, which will stop modeling if the loss is not further reduced in the last 3 iterations. Results show that convergence is typically achieved within 3 iterations for this dataset, and increasing the maximum number of iterations beyond this does not further change the results. Notably, even a single iteration already yields competitive performance, likely benefiting from the strong initialization based on unspliced-to-spliced temporal delay.

Overall, these results highlight a trade-off between computational efficiency and modeling expressiveness. While TSvelo is more computationally demanding than classical approaches, it provides a more flexible framework for incorporating regulatory information and capturing complex gene interactions, which we believe justifies the additional computational cost in scenarios requiring accurate dynamical inference.

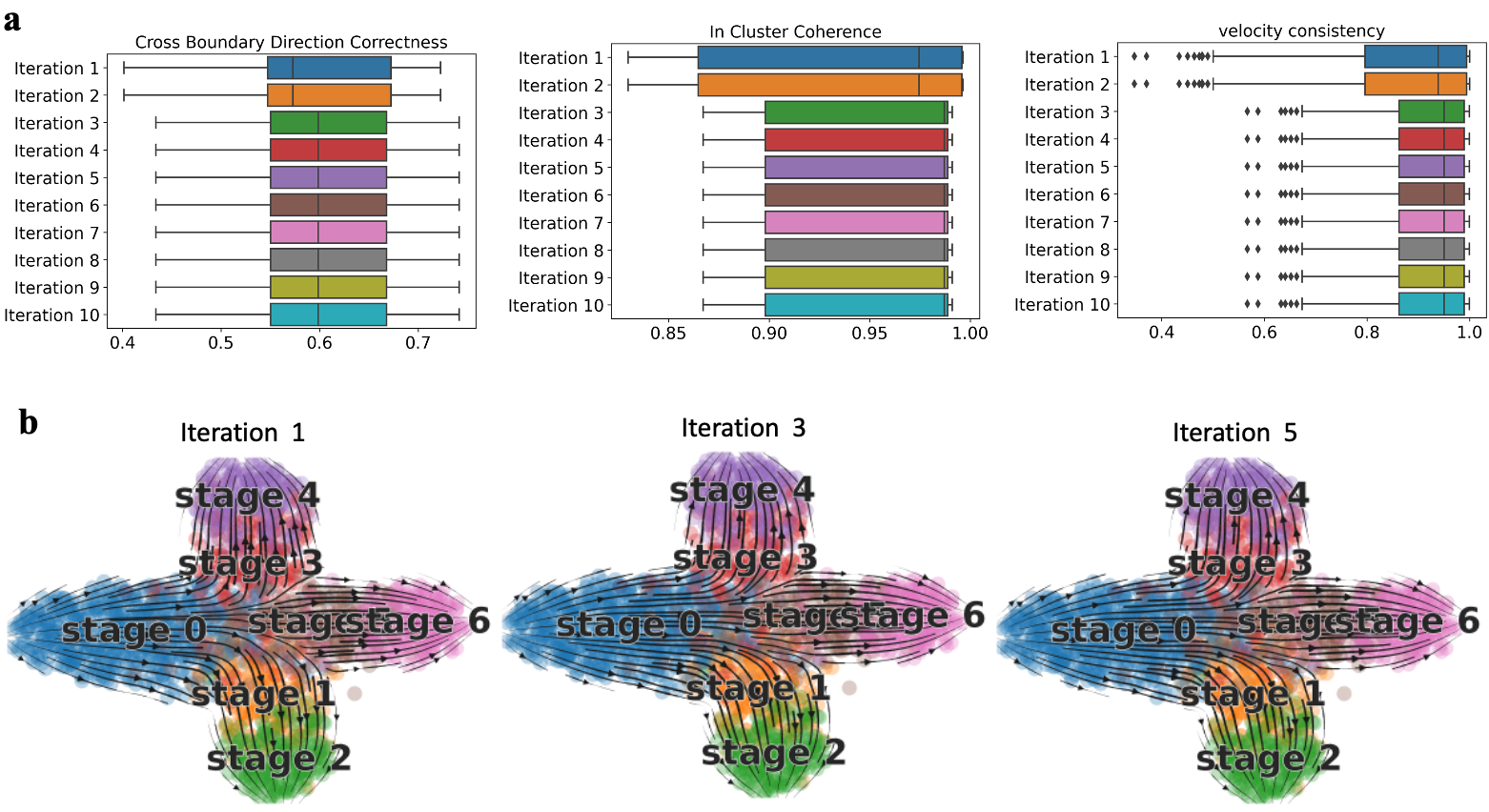

**Figure S16. Effect of the number of EM iterations on TSvelo performance in the simulation dataset.** (a) Quantitative evaluation across different iteration settings. (b) 2D visualization of inferred dynamics. The results show that TSvelo converges rapidly, and performance remains stable after 3 iterations due to the built-in early stopping mechanism in the EM framework.

**Table S3. Runtime of TSvelo under different numbers of EM iterations on the simulation dataset.** The results demonstrate that TSvelo achieves stable performance even with a small number of iterations. Due to the early stopping mechanism of the EM framework, since TSvelo get the best results at iteration 3, it actually performs no more than 6 iterations.

| **Number of max iterations set in hyperparameters** | 1 | 2 | 3 | 4 | 5 |
| --- | --- | --- | --- | --- | --- |
| **Running Time (Sec)** | 1007 | 1168 | 2145 | 2208 | 2989 |
| **Number of max iterations set in hyperparameters** | 6 | 7 | 8 | 9 | 10 |
| **Running Time (Sec)** | 3120 | 3256 | 3126 | 3121 | 3027 |

**Null simulations for evaluating RNA velocity approaches on data without underlying dynamic structure**

We have added null simulations to evaluate TSvelo and baseline approaches on data without underlying dynamic structure. Specifically, we generated a null dataset including 200 genes and 600 cells by independently sampling spliced (S) and unspliced (U) counts, thereby removing any coherent transcriptional relationship between them.

When applying scVelo and UniTVelo to this data, no genes passed the velocity gene selection step under the default likelihood-based filtering, and no velocity field could be obtained. We further tested TSvelo, Dynamo, and cellDancer on the same null data and observed that all three methods still produce trajectory-like patterns despite the absence of true dynamics (**Fig. S17**).

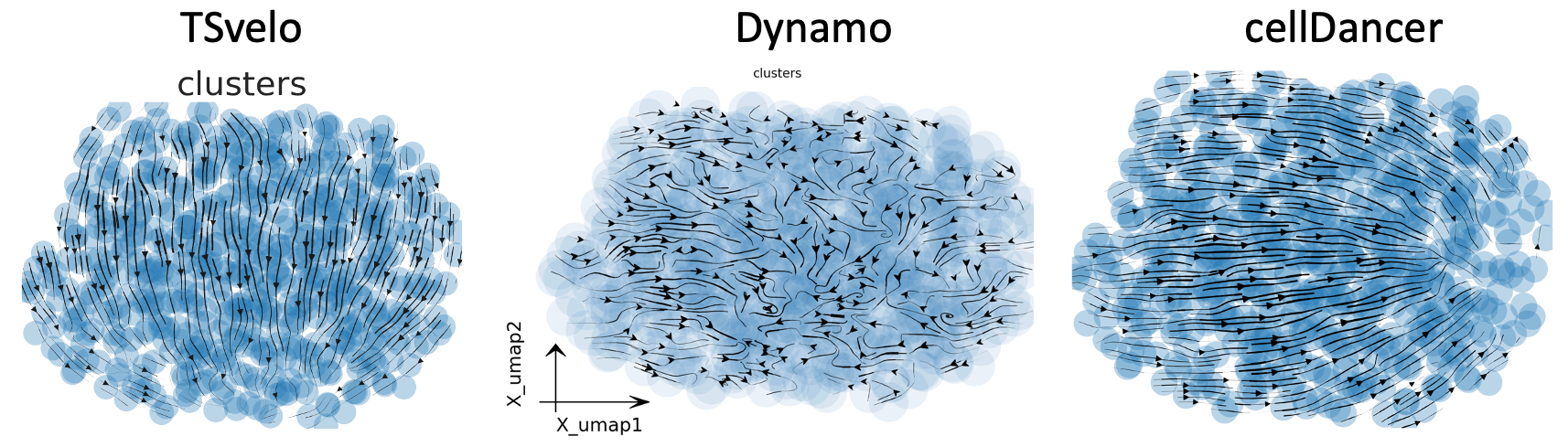

**Fig. S17. The predicted velocity streams of TSvelo, Dynamo, and cellDancer on the null simulation data.**

Including TSvelo, many RNA velocity and trajectory inference approaches assume that they are applied to datasets reflecting underlying dynamic biological processes. Incorporating additional checks during preprocessing could help prevent applying velocity analysis to non-dynamic datasets.
